# Thermodynamically controlled biokinetics modelling of a lindane-degrading anaerobic enrichment culture

**DOI:** 10.64898/2026.10.01.755988

**Authors:** Jingzhi Liang, Line Lomheim, Luz A. Puentes Jácome, Elizabeth A. Edwards, Brent E. Sleep

## Abstract

The anaerobic biotransformation of lindane (γ−hexachlorocyclohexane), a persistent legacy pesticide and challenging subsurface contaminant was modelled using an integrated Monod biokinetics framework. Calibrated against batch experimental data, this model accounted for the dissolution of solid phase lindane and the thermodynamics of reactions involved. The study focused on a methanogenic, lindane-degrading enrichment culture (GT2) that used ethanol as the electron donor, where ethanol fermentation supplied hydrogen to support reductive dechlorination of lindane. A *Dehalobacter* strain was identified as the lindane-degrading organism, while a member of bacterial order Geobacterales carried out ethanol fermentation. First-order decay constants for lindane-degrading *Dehalobacter* were estimated from experimental data and showed variation across different growth conditions. The biokinetics model captured interactions among key microbial processes, namely lindane dechlorination, ethanol fermentation, hydrogenotrophic methanogenesis, and acetoclastic methanogenesis. Thirteen biokinetics parameters including maximum substrate utilization rates (*k*), half-velocity constants (*K_s_*), and decay constants for these microbial processes were estimated through a Bayesian optimization approach. Global sensitivity analysis showed that eight parameters, primarily *k* and *K_s_*, had significant linear impact on the model output, whereas responses to decay constants were more variable and potentially non-linear. This study proposed an effective modelling approach for characterizing syntrophic microbial growth within enrichment cultures and provides valuable biokinetics parameters estimates for real-world applications of anaerobic lindane bioremediation.

**Synopsis:** Integrated biokinetics modelling of syntrophic lindane degradation and microbial interactions, incorporating decay constant estimation to inform anaerobic bioremediation design decisions.

**Graphical abstract:** 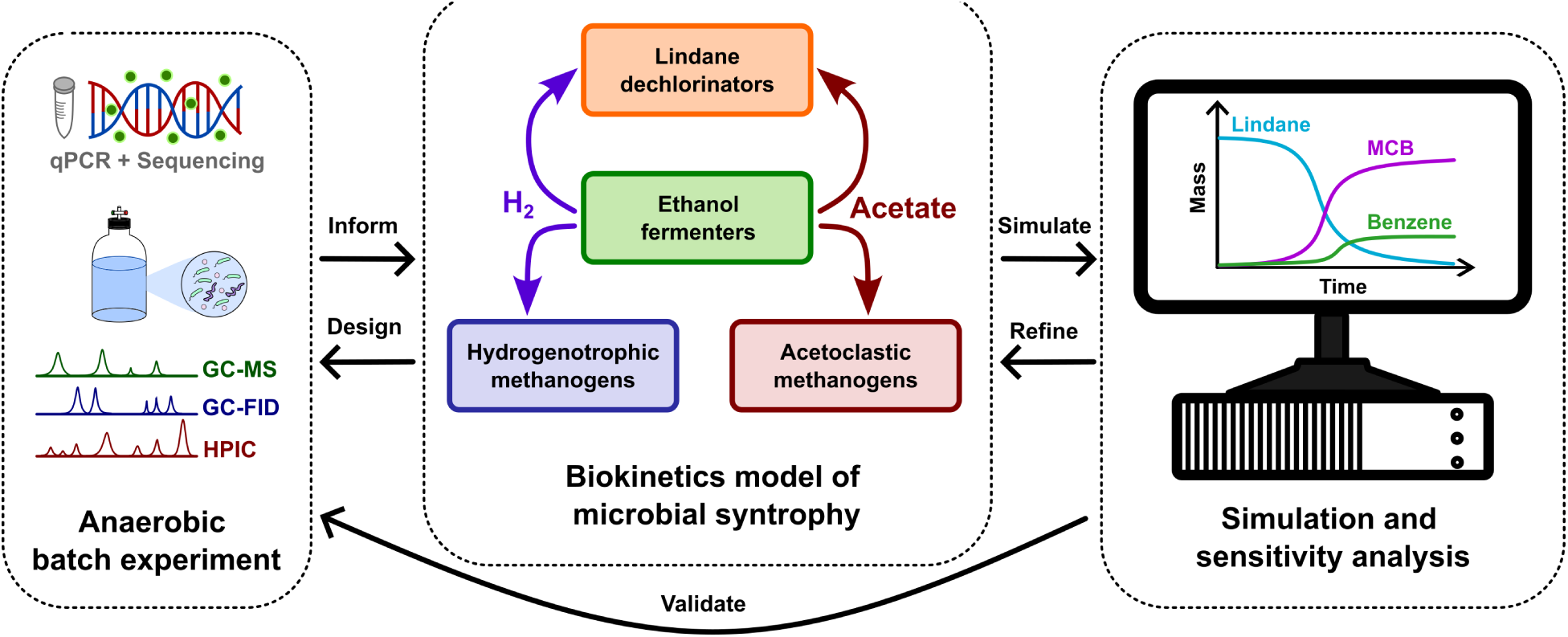

## 1. Introduction

Lindane, the γ isomer of hexachlorocyclohexane (HCH), was widely used as a broad-spectrum chlorinated pesticide and fungicide since 1940s.^1, 2^ Worldwide applications of this insecticide contributed to agricultural yield, livestock protection, and the prevention and treatment of vector-transmitted diseases in humans. Up to 10 million tons of lindane were released into the environment during its half-century of usage before it was banned in most countries globally by late 1990s.^3^ The insecticidal property of lindane results from its ability to reduce neurotransmission inhibition, leading to hyperexcitation and death upon acute high-dosage exposure.^4^ Meanwhile, adverse health impacts of chronic low-dosage exposure to lindane include probable carcinogenicity, immunotoxicity, and genotoxicity.^5^ Owing to its high lipophilicity (log K_ow_ = 3.83, water solubility = 7.2 mg/L at 25 ℃),^6^ lindane can easily accumulate inside of the organism that is exposed to it, subsequently biomagnifying and bioaccumulating through the food chain. Its high lipophilicity also explains its preferential partition into organic materials in the sediment phase, resulting in its low mobility potential and high adsorption rate in soil. In spite of this, monitoring studies show that lindane can be released into groundwater - one of the two major sources of drinking water worldwide - through soil leachate.^7–9^ To date, this toxic compound has been found in all environmental media in common aquatic and terrestrial organisms worldwide.^10^ Given its significant toxicity to human health and the environment and its potential to persist and transport far through atmospheric circulation, the Stockholm Convention on Persistent Organic Pollutants listed lindane to the list for elimination in 2007.^11^

To remove lindane from the subsurface, in situ bioremediation involving microorganisms capable of transforming lindane is considered more cost-effective with minimal disruption to the local environment compared to conventional ex situ approaches. In the anaerobic environment that prevails underground, microbial reductive dechlorination of lindane yields a mixture of monochlorobenzene (MCB) and benzene as stable end products.^12–18^ Anaerobic microorganisms belonging to genera *Clostridium*,^19–25^ *Desulfococcus*,^17^ *Desulfovibrio*,^17^ *Dehalococcoides*,^18^ and *Dehalobacter*^12, 13^ are known to carry out the reductive biotransformation of lindane. Although the anaerobic degradation of lindane was reported as early as 1966,^26^ current field applications of lindane bioremediation mostly rely on its aerobic degradation pathway.^27–29^ The anaerobic pathway has been overlooked primarily because it terminates at MCB and benzene, both of which are still toxic in the environment, making the anaerobic detoxification of lindane particularly challenging. With the discovery of anaerobic bioremediation processes for MCB and benzene,^30, 31^ the anaerobic sequential degradation of lindane to non-toxic end products has been proven to be possible.^12^ However, the kinetic parameters describing the degradation scheme proposed in these studies have yet to be explored and investigated to inform future engineering design.. This study aims to bridge the gap between empirical work and the engineering application of anaerobic lindane biotransformation through numerical modelling.

Microbial kinetic modeling is the mathematical characterization of the active microbial system and its physicochemical interactions with various environmental factors, which include reaction substrates and their concentrations, pH, temperature, pressure, and mass transfer between different phases. It connects the active biomass and substrates systematically and quantitatively and is therefore considered an essential tool for engineering design and operation.^32^ However, published datasets on anaerobic lindane degradation are often inadequate due to low sampling frequency during a feeding cycle and/or the lack of direct quantification of lindane-degrading microorganisms,^13, 15–18, 33^ which limits the accurate estimation of kinetics parameters commonly used to predict remedial performance in the field. This highlights the need for comprehensive biokinetics study herein.

In this study, a batch experiment was set up using a well-maintained lindane-degrading enrichment culture, providing essential data for model calibration and validation. Molecular biology tools were used to monitor temporal variations in the microbial community structure and biomass concentration of major functional microbial groups in different growth phases and conditions. The mass transfer between the aqueous and solid phases of lindane was also accounted for, mimicking the slow release of the compound in the environment. Since anaerobic microbial processes often operate close to thermodynamic equilibrium, Gibbs free energy calculations were incorporated into the modelling framework to prevent thermodynamically unfavorable conversions. This approach is consistent with previous studies emphasizing the importance of reaction thermodynamics in modelling anaerobic and syntrophic microbial processes.^34–36^ The modelling approach also considered minimum cell energy requirement, biomass decay, and the thermodynamics of metabolic and anabolic reactions of syntrophic microbial groups for growth yield estimations. Through detailed observation and frequent measurements of key components in the reaction system, a comprehensive biokinetics model was established using MATLAB^®^ Simulink^®^. Results from this study have enabled us to: (1) disentangle the inter-relationships between different microbial groups within the culture; (2) derive a set of biokinetics parameters for these microbial groups, especially for the anaerobic lindane-degrading microorganisms; and (3) pave the way for future biokinetics studies on anaerobic lindane biotransformation.

## 2. Materials and methods

### 2.1. Chemicals

All chemicals used in this study were purchased through Sigma-Aldrich (Canada). Lindane was obtained as PESTANAL^®^ analytical standard (purity ≥ 98.0%) as powder. HPLC-grade ethanol and acetone as well as MilliQ water were used for preparing stock solutions. Gases for maintaining anaerobic culturing were supplied by Praxair (Canada).

### 2.2. Enrichment culture and experimental setup

The lindane-degrading culture (GT2) modelled in this study was originally enriched from a sample of contaminated agricultural lands collected in Guadeloupe (French West Indies) in 2010.^12, 13^ Previous studies in our group investigated how the microbial community was impacted by various HCH isomers.^13^ The culture is maintained in a pre-reduced defined anaerobic growth medium,^37^ which uses sodium bicarbonate as a pH buffer to sustain microbial growth under near-neutral pH conditions. *Dehalobacter* (*Dhb*) in this culture was identified as the lindane degrader, relying on a syntrophic mechanism where hydrogen required for reductive lindane dechlorination was supplied through ethanol fermentation, while both acetoclastic and hydrogenotrophic methanogens (e.g., *Methanothrix*, *Methanoregula*) were found in GT2 and may compete with *Dhb* for hydrogen and carbon sources such as acetate.^12, 13^ The target lindane concentration was 100 μM culture with ethanol amended in excess at a 5 to 2 molar ratio to lindane. An average lindane degradation rate of 8.1 μM/day was determined prior to the batch experiment. The products of MCB dechlorination, benzene and MCB, were produced at a molar ratio that ranged from 0.05 to 0.11.

The batch experiment to determine kinetic constants was set up using 250 mL sterile serum bottles with mininert caps in four groups: the experimental (EXP) group, the *Dhb* starved control (DSC) group, the starved control (SC) group, and the analytical control (AC) group (Table S1). Bottles in the first three groups were prepared in quadruplicate and inoculated with 20% (v/v) GT2 culture. The EXP bottles received lindane and ethanol, DSC bottles were fed with ethanol only, and the SC bottles were not fed any external electron donor or acceptor. AC group bottles were prepared in triplicates using the sterile growth medium without culture inoculation. The same lindane and ethanol dosages as the EXP bottles were amended to the AC bottles. The predicted maximum amount of benzene, MCB, and methane in the biotransformation products were added to the AC bottles to control for possible instrumental drifts and abiotic lindane degradation during the experiment. All bottles were filled with a total liquid volume of 220 mL.

Amending bottles with lindane presents a challenge because of its low aqueous solubility (7.2 mg/L at 25°C^6^). A concentrated stock (47 mg/mL) was first prepared in acetone. An aliquot (136 μL) of this acetone stock was added into each empty 250 mL bottle and acetone was evaporated to dryness under a stream of N_2_ while the bottle was swirled, leaving 20 μmol solid lindane per bottle. All bottles, including lindane-preloaded EXP and AC bottles, were capped using aluminum foil and then moved into an anaerobic chamber (Coy Lab Products) supplied with CO_2_/H_2_/N_2_ (10%/10%/80%) and pure nitrogen for anaerobic stabilization for a minimum of 2 days prior to subsequent bottle set-up, inoculation, feeding, incubation, and sampling procedures. The anaerobic chamber atmosphere was maintained at approximately 1% H_2_ throughout the experiment. On the inoculation day (Day 0), the GT2 inoculum was purged with a CO_2_/N_2_ (20%/80%) gas mix for 40 minutes to remove any residual volatile compounds that had accumulated during culture maintenance. Sterile pipettes were then used to transfer sterile medium and purged culture inoculum directly into each bottle according to the treatment plan (Table S1). For bottles receiving ethanol in the EXP and DSC groups, 6.42 μL of 50% (v/v) aqueous ethanol solution (providing 55 μmol ethanol) was injected through the mininert cap. Two milliliters of the headspace gas in the AC bottles were replaced with pure methane gas (Restek). To avoid possible disturbance to the culture biokinetics induced by volume changes upon liquid sampling,^38^ sampling volumes were as small as possible such that the final liquid volume was ∼190 mL (∼86% of the original). Moreover, to ensure cross-comparison among groups, the same volume of liquid was removed across all bottles on each sampling day.

### 2.3. Analytical methods

Detailed descriptions of analytical methods for measuring lindane, MCB, benzene, methane, and volatile fatty acids (VFAs) can be found in Text S1, Text S2, and Text S3 in Supplementary Information, A summary of each method is listed below. Lindane concentration was quantified using a gas chromatograph equipped with a mass spectrometry (GC-MS) system (Agilent 7890B and 5977B GC/MSD). Briefly, 500 μL of aqueous sample was added into 9.5 mL of HCl-acidified NaCl solution (10%, wt/vol, in Milli-Q water, pH = 2.5) for enhanced lindane extraction efficiency by solid phase microextraction (SPME). The acidified NaCl solution also quenched microbial activities after sampling. Concentrations of benzene, MCB, and methane were measured by a GC (Agilent 7890A) with a flame ionization detector (FID). For this analysis, 0.5 or 1 mL sample liquid was added to 5 mL of acidified water (prepared using 6N HCl and MilliQ water, pH <2) to stop further microbial activities. Concentrations of VFAs (acetate and propionate) were quantified using a Dionex Integrion High-Pressure Ion Chromatography (HPIC) system (Thermo Scientific) for anions. Prior to this analysis, each sample (minimum 500 μL) was filtered using a 0.2 μm filter into a 1.5 mL micro-centrifuge tube and stored at -20°C. Due to instrumental limitations, the hydrogen concentration was not directly measured in this study.

### 2.4. Molecular methods

DNA was purified from 2 mL samples of culture liquid. Each sample was centrifuged for 20 minutes at 13,000 rpm at 10 ℃ to obtain a cell pellet for semi-automated DNA extraction by KingFisher Duo Prime (Thermo Fisher Scientific) and a MagMax Microbiome DNA extraction kit (Thermo Fisher Scientific). The resulting purified DNA was subsequently quantified using an Invitrogen Qubit 3.0 Fluorometer (Thermo Fisher Scientific). Quantitative polymerase chain reaction (qPCR) assays were carried out to measure 16S rRNA gene copy concentrations of *Dhb*, bacteria and archaea using their respective DNA primers (Table S2).

A total of 57 DNA samples were sent for 16S rRNA gene amplicon sequencing at Genome Quebec Innovation Centre (McGill University), where the DNA was checked for quality and sequenced using the Illumina MiSeq PE300 platform. A modified primer set with “staggered ends” of 926f and 1392r (926f-modified: AAACTYAAAKGAATWGRCGG; 1392r-modified: ACGGGCGGTGWGTRC) targeting the V6-V8 region of the 16S rRNA gene of bacteria and archaea as well as the 18S rRNA gene of eukarya was used for improved sequencing quality.^39^ Raw reads were pre-processed using the DADA2 pipeline of QIIME 2 version 2022.2.^40^ Chimeric sequences and sequences shorter than 400 bp were excluded from further processing. The amplicon sequence variants (ASVs) were classified using the SSU SILVA 132 database^41^ version 2022.6 trained against the modified primer set. Further processing and analyses were performed in Excel. All singleton ASVs (those that appeared only once in all samples) and contaminant ASVs (those detected in the sequencing blank sample and were in very low abundance in other samples) were removed. Raw sequencing data from this study have been deposited in the NCBI Sequence Read Archive under BioProject PRJNA1143471.

### 2.5. Modelling approach

Biokinetics modelling was developed based on the primary assumption of mass balance, accounting for changes in liquid volume and mass removal due to liquid sampling. The model employed *Monod* kinetics and incorporated thermodynamic equilibria in anaerobic biochemical reactions. A general model workflow is presented in Figure 1, where starting concentrations of chemicals and biomasses are defined during system initialization. Multiplicative *Monod* kinetics was applied to model lindane dechlorination, describing the effect of H_2_ limitation.^42^ In the biokinetics block, four functional microbial processes (lindane dechlorination, ethanol fermentation, hydrogenotrophic methanogenesis, and acetoclastic methanogenesis) were modelled separately (Table 1). For lindane dechlorination, the stoichiometric ratio between MCB and benzene (15:1) was derived from experimental data of this study. The revised thermodynamic electron equivalents model (TEEM2) was adopted in the thermodynamics block to predict microbial yields (Text S10).^43^ To simulate biomass formation, a typical empirical cell biomass formulation of C_5_H_7_O_2_N was used, and a universal energy transfer efficiency of 0.6 was applied, as recommended for anaerobic microorganisms and those growing in mixed cultures.^44, 45^ The use of a generic biomass formula provides a consistent and widely accepted basis for stoichiometric and thermodynamic calculations, given that cellular composition can vary across taxa and growth conditions in microbial consortia such as GT2, and culture-specific measurements were not available. Acetyl-CoA was used as the intermediary metabolite for *ΔG* calculations for anabolism as suggested in TEEM2.^43^ Ethanol was assumed to be the sole carbon source for ethanol fermenters, while acetate was assumed to be the carbon source for lindane degraders and all methanogens.^46^ Although hydrogenotrophic methanogens can use inorganic carbon (e.g., CO_2_) as a carbon source, acetate remains an important growth factor for biosynthesis.^47–50^ While TEEM provides an approach for estimating autotrophic biomass yields using inorganic carbon as the carbon source, its formulation relies on a water-oxygen half-reaction as a thermodynamic reference and was therefore not applied to the anaerobic autotrophic methanogens considered here.^43^ Acetate was selected as the carbon source for all methanogens for simplicity. For all cell synthesis reactions, ammonium was assumed as the nitrogen source. Concentrations of key compounds affecting reaction thermodynamics were modelled in the simulation, including lindane, MCB, benzene, acetate, hydrogen, methane, and chloride. Among these key compounds, hydrogen was the only one not directly measured during the experiment. Its initial concentration was instead inferred based on the atmospheric composition inside the anaerobic chamber (i.e., 1% H_2_). Using Henry’s Law, the partitioning between aqueous and gaseous phases was considered for volatile compounds hydrogen, MCB, benzene, and methane (Table S8), without considering salinity or temperature effects. The mass transfer of lindane from solid to aqueous phase was modelled through first-order kinetics^51, 52^ (Equation 1) where a lumped mass-transfer rate coefficient (*K_l_*) was estimated from lindane measurements in the AC bottles (Text S7):

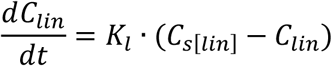

**Equation 1. First-order lindane dissolution kinetics, where *C_lin_* represents aqueous concentration, *C_s[lin]_* is solubility limit, and *K_l_* is the lumped mass-transfer rate coefficient.**

**Figure 1.**
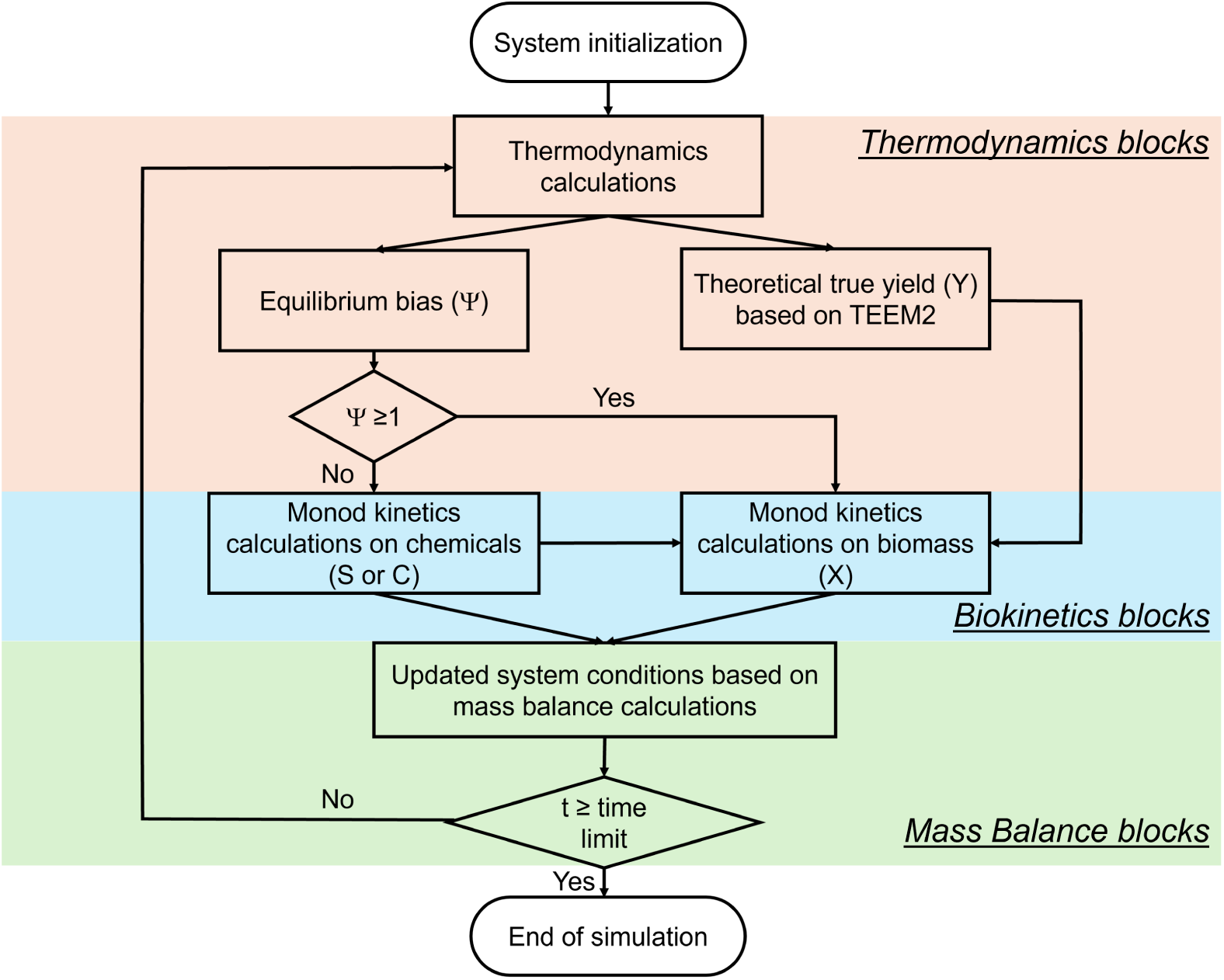
General workflow of the GT2 biokinetics model.

**Table 1.** Stoichiometry and Gibbs free energies of reactions in the GT2 biokinetics (25℃, pH = 7).

| Microbial Group | Energy Reaction Equation | $\Delta G^{\circ\prime}$ (kJ/mol e<sup>-</sup>eq) | # e<sup>-</sup><br>eq/reaction |
| --- | --- | --- | --- |
| Lindane dechlorination | $16\text{C}_6\text{H}_6\text{Cl}_6 + 33\text{H}_2 = \text{C}_6\text{H}_6 +$<br>$15\text{C}_6\text{H}_5\text{Cl} + 81\text{HCl}$ | -24.03 | 450 |
| EtOH fermentation | $\text{C}_2\text{H}_5\text{OH} + \text{H}_2\text{O} = 2\text{H}_2 + \text{H}^+$<br>$+ \text{CH}_3\text{COO}^-$ | 0.80 | 12 |
| H<sub>2</sub> methanogenesis | $4\text{H}_2 + \text{HCO}_3^- + \text{H}^+ = \text{CH}_4 + 3\text{H}_2\text{O}$ | -16.95 | 8 |
| AC methanogenesis | $\text{CH}_3\text{COO}^- + \text{H}_2\text{O} = \text{CH}_4 + \text{HCO}_3^-$ | -3.88 | 8 |

### 2.6. Thermodynamically controlled biokinetics

Using previously published standard Gibbs free energies of formation (Table S5), Gibbs free energy of reactions involved in thermodynamic calculations were derived from organic half-reactions of electron donors or acceptors (Table S6). For each half-reaction, the Gibbs free energy was calculated assuming pH = 7 and temperature at 25℃, with the bicarbonate ion as the oxidized form of carbon placed on the left side of the equation, and the reduced compound and water on the right (Table S6).^44, 53^ To derive the actual Gibbs free energies during model simulation, the initial concentrations of bicarbonate and chloride were calculated based on the medium recipe as 12.26 mM and 10.88 mM, respectively. The initial concentration of hydrogen was estimated based on the partial pressure of hydrogen (1%) in the anaerobic chamber. The initial acetate concentration was estimated based on the IC measurement on Day 0 of the SC bottles of 10 μM (Table 13). For cell synthesis reactions, the intracellular concentration of acetyl-CoA was estimated at 310 μM, an average value measured in *Escherichia coli* cells,^54^ and remained constant throughout the simulation. For simplicity, the activity coefficients of all reactants and products were set to one, and their activities were approximated by their molar concentrations since the system is dilute, operated at room temperature, maintained a neutral pH, and was kept at 25 °C. Critical Gibbs free energy (Δ*G*_cr_) was incorporated into the model to describe the physiological requirements of microorganisms prior to ATP production.^55^ It represents the minimal energy needed for microbial survival and is related to the activation energy for all enzymatic reactions within microbial cells.^42^ Therefore, the net Gibbs free energy change available for microbial growth is *θ* = Δ*G*_e_ - Δ*G*_cr_, where Δ*G*_e_ is the Gibbs free energy of metabolic energy reactions. A negative *θ* value allows extra energy for ATP formation and cell synthesis. Δ*G*_cr_ was set as -1 kJ per mole electron equivalent (e^-^eq) for each energy reaction and remained constant. This value was estimated from previous empirical studies on the bioenergetics of syntrophic-methanogenic cocultures, where Δ*G*_cr_ ranged from -3.8 to -18.2 kJ per mole of donor reaction for butyrate or ethanol oxidation, each transferring 4 e^-^eq.^56, 57^ During model simulations, Δ*G*_e_ is calculated using instantaneous concentrations of substrates and products for each metabolic reaction.

To capture the reversible nature of enzymatic biochemical reactions^58^ under anaerobic conditions where the degradation rate of electron acceptors is limited by the threshold concentration of the electron donor and end-product inhibition,^34, 59–62^ a thermodynamic term (*Ψ*) was introduced into the biokinetics model, describing the instantaneous “deviation to equilibrium” of each metabolic reaction. In this model, thermodynamic feedback was characterized by extending a reverse kinetic model,^61^ considering the net Gibbs free energy change (*θ*), and adapting it for Monod kinetics.^63^ In the modelled system, reaction equilibrium is reached when *θ* = 0, resulting in net zero substrate consumption. *K*_eq_ represents the reaction quotient under equilibrium conditions when Δ*G*_e_ equals Δ*G*_cr_. By definition, *Ψ* = *Q*/ *K*_eq_, where *Q* is the instantaneous reaction quotient. *Ψ* can also be expressed as (Equation 2):

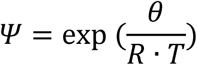

**Equation 2. Equilibrium bias (*Ψ*) for enzymatic reversible reactions, where *θ* is the net Gibbs free energy (kJ/mol) for cell growth at temperature *T* (K), and the ideal gas constant *R* is expressed in kJ/mol/K.**

The incorporation of the thermodynamic term into the biokinetics model enables more realistic simulation of microbial processes in the anaerobic environment that operate close to equilibrium condition.^62^ For metabolic reactions with only one rate-limiting substrate and one rate-limiting product, as for ethanol fermentation and hydrogenotrophic/acetoclastic methanogenesis, a simple modification to the Monod kinetics was applied (Equation 3), and that substrate consumption would cease when *Ψ* equals to 1:

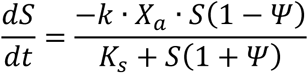

**Equation 3. Biokinetic model for ethanol fermentation and methanogenesis, where *S* is substrate concentration (μM), *k* is maximum specific substrate utilization rate (μmol/mg/day), *X_a_* is active biomass concentration (mg/L), *K_S_* is half-velocity constant (μM), and *Ψ* is equilibrium bias (unitless).**

For anaerobic lindane dechlorination, a similar modification was applied to the multiplicative Monod kinetics (Equation 4). Though both lindane concentration and hydrogen concentration can be rate limiting, the (1+*Ψ*) term was applied to the hydrogen concentration term. Hydrogen, the electron donor for both dechlorination and hydrogenotrophic methanogenesis, has been reported as the primary rate-limiting factor for reductive dechlorination in empirical studies of methanogenic cultures.^42, 64, 65^

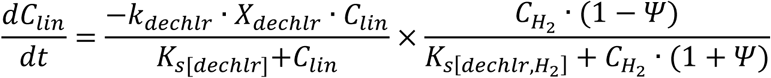

**Equation 4. Biokinetic model for anaerobic lindane dechlorination, where *C_lin_* is lindane concentration (μM), *k_dechlr_* is maximum specific substrate utilization rate for lindane dechlorination (μmol/mg/day), *X_a_* is active biomass concentration (mg/L), *K_S_*_[*dechlr*]_ is half-velocity constant for lindane during dechlorination (μM), *C_H2_* is hydrogen concentration (μM), *K_S_*_[*dechlr, H2*]_ is half-velocity constant for hydrogen in lindane dechlorination (μM) and *Ψ* is equilibrium bias (μM).**

### 2.7. Parameter estimation and sensitivity analysis

The decay coefficients for *Dhb* and ethanol fermenters were independently estimated from experimental data assuming first-order decay using the Nonlinear Curve Fit Tool of OriginPro (2023b), see Text B9 for details. Estimation of 13 biokinetic parameters including specific substrate utilization rates, half-velocity constants, and decay constants for all four microbial groups was conducted by interacting with the Simulink model using a MATLAB script (R2024a). During each parameter estimation run, measured time series data from bottle EXP2 (Table S13) were used as calibration data and compared with simulation results. The calibration dataset included eight expected model outputs: the masses of lindane, MCB, benzene, and methane, as well as the biomass concentrations of *Dhb*, EtOH fermenter, acetoclastic methanogens, and hydrogenotrophic methanogens. Biomass concentrations were determined by qPCR for *Dhb*, or by a combination of qPCR and 16S rRNA sequencing data for EtOH fermenter, acetoclastic methanogens, and hydrogenotrophic methanogens, as described in Text S5, and were converted to mg/L using the conversion factors introduced in Text S6. Bottle EXP2 was selected because it provided a complete calibration dataset for lindane, MCB, benzene, methane, and microbial biomass dynamics. Data from the other experimental bottles were used to evaluate replicate variability and support interpretation of the observed trends, but were not included in parameter estimation. The initial state of the model was set up using data collected on Day 0 in EXP2. Initial guesses and boundaries for estimated parameters are provided in Table S10. The optimized fitting of measured and simulated outputs ensured minimized cost function (weighted sum of squared errors, WSSE, Equation 5) through Bayesian optimization (BO) (MATLAB; bayesopt) using the acquisition function of expected-improvement-plus within 200 iterations (convergence threshold = 10^-9^). In Equation 5, the first four outputs are from chemical data and the last four outputs are biomass data, where *w*_1_ = 0.8 is the assigned weight for the former, *w*_2_ = 0.2 is the assigned weighted for the latter, *r_i, j_* is the residual for *i*th time point of the *j*th output normalized using the maximum value of the calibration data for output *j*, and *N_j_* is the total number of time points for output *j*. The use of weighting in parameter estimation is intended to prioritize the fitting of chemical measurements, which generally have smaller relative errors than biomass measurements. One full dataset from EXP2 and 100 datasets generated through leave-one-out-bootstrapping (LOOB) method through randomly dropping one time point in each target output were used in BO to calculate the statistics of estimated parameters.

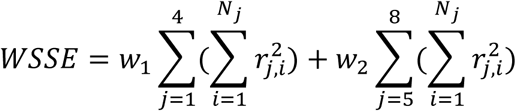

**Equation 5. Definition of weighted sum of squared errors for parameter estimation for the GT2 biokinetics model.**

To quantify the relative contribution of model parameters to simulation results, global sensitivity analysis (GSA) was carried out on the 13 parameters estimated using this model under the EXP2 scenario. One thousand sets of parameters were randomly generated through Latin hypercube sampling,^66^ a stratified sampling method designed to provide efficient coverage of the parameter space, assuming uniform distribution within the boundaries specified in Table S10. The impact of choices of parameter values was then assessed through signal matching between simulated and measured data for both the global output (WSSE) as in parameter estimation as well as individual output signals (sum of squared normalized errors, SSE). Four complementarty approaches were applied in GSA because they characterize different types of parameter-output relationships.Linear impacts were analyzed through standardized regression and partial correlation, while monotonic responses were assessed through Spearman rank correlation. Morris screening was additionally used to identify parameter effects that maybe nonlinear or dependent on interactions with other paramters.^67^ Significance testing was conducted for the first three approaches. In the Morris method, 52 trajectories were generated (step size = 0.1) to evaluate the elementary effects (EE) of each parameter. The mean absolute elementary effect (|EE|) was used as an indicator of the overall influence of a parameter on the model output, while the standard deviation (SD) of the EEs characterized the variability of this influence across the parameter space. A larger SD may indicate nonlinear effects and/or interactions with other parameters. Unlike the coefficients obtained from the other three approaches, the Morris method generates multiple EEs for each parameter; therefore, the SD of these EEs is reported as a separate metric. These sensitivity measures were interpreted as complementary indicators of parameter influence rather than definitive evidence of sensitivity.

## 3. Results and Discussion

### 3.1. Syntrophic microbial community of GT2

In the batch experiment, approximately 600 mL of GT2 culture liquid, prepared by thoroughly mixing three bottle of the GT2 culture, were used to inoculate the EXP, DSC, and SC groups. The inoculum culture was sampled in duplicate for DNA analysis, and both DNA samples exhibited nearly identical microbial community structures at the ASV level (Figure S5), indicating good mixing results and allowing for cross-comparison between inoculated bottles. In the GT2 culture, Geobacterales_(o)_ASV1 was the most abundant bacterial group, making up 35 ± 2% of all bacterial taxa, followed by Geobacteraceae_(f)_ASV1 (17 ± 1%), *Sulfurospirillum*_ASV1 (9 ± 1%), and *Dehalobacter*_ASV1 (6 ± 2%) (Figure S5: A). Within the domain Archaea, acetoclastic *Methanothrix*_ASV1 and *Methanothrix*_ASV2, and hydrogenotrophic *Methanoregula*_ASV1 were dominant, accounting for 49 ± 3%, 11 ± 0.2%, and 15 ± 2% of all archaeal taxa, respectively (Figure S5: B).

A gradual enrichment of *Dehalobacter* was observed in the EXP group, which was fed with lindane and inoculated with GT2, between Day 0 and Day 27 (Figure 2: A). Consistent with previous studies identifying *Dhb* as the lindane dechlorinator^12, 13^, the relative abundance of *Dhb* within the total bacterial community, as determined through qPCR, increased from 6 ± 2% on Day 0 to 22 ± 6% on Day 27. This relative abundance then dropped to 4 ± 1% by Day 117, when lindane was completely depleted in all EXP bottles (Table S11). In the DSC and SC groups, where lindane was not amended, the relative abundance of *Dhb* decreased throughout the monitoring period, indicating the decay of the dechlorinator due to starvation, in this case, lack of electron acceptor (Figure 2: A).

**Figure 2.**
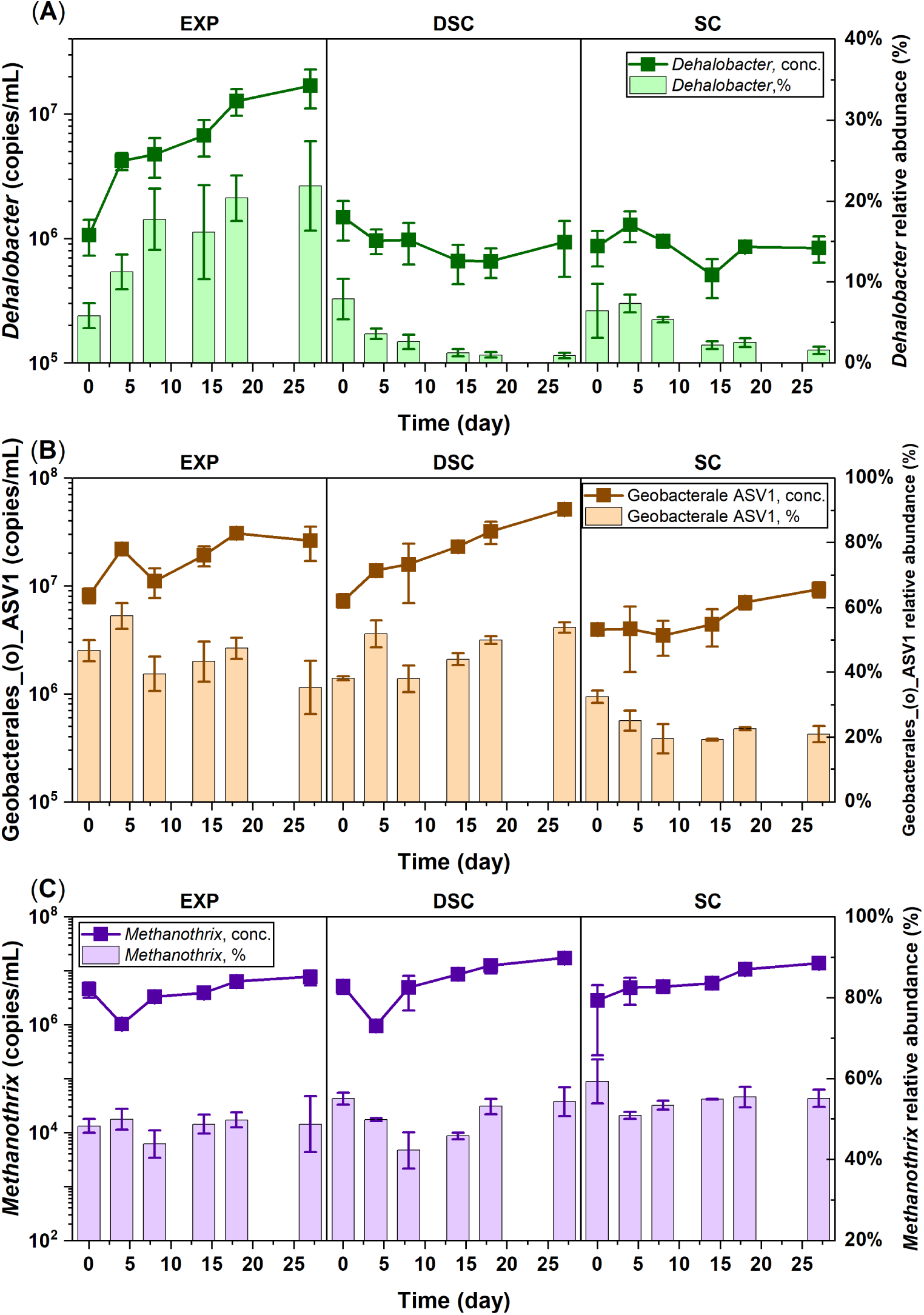
Absolute and relative abundances of key microbial groups during the first 27 days of the monitoring period in the experimental (EXP), *Dehalobacter* starved control (DSC) group, and starved control (SC) groups: (A) *Dehalobacter*, (B) Geobacterales_(o)_ASV1, and (C) *Methanothrix*.

In the DSC group was amended only with ethanol, an evident enrichment of Geobacterales_(o)_ASV1 was observed between Day 0 and Day 27, with its relative abundance (as a percentage of total bacteria) increasing from 38 ± 1% to 54 ± 2% (Figure 2: B). In contrast, relative abundance of Geobacterales_(o)_ASV1 in the SC group, which did not receive any substrates, declined from 32 ± 2% on Day 0 to 21 ± 3% on Day 27 (Figure 2: B). Therefore, this bacterial strain was hypothesized to be the major ethanol fermenter in the GT2 culture, producing hydrogen and the carbon source, acetate, for lindane dechlorination by *Dhb* and methanogenesis. Hydrogenotrophic acetogenesis may also occur in the culture due to the presence of *Acetobacterium*_ASV1 (Figure S3: A). However, this *Acetobacterium* strain comprised 2 ± 0.2% of the total bacterial community in the inoculum culture based on ASV relative abundance and was therefore much less prominent as a potential contributor to acetate production than the major ethanol fermenter.

For groups that received ethanol (EXP and DSC), a clear enrichment of the acetoclastic methanogen *Methanothrix* was observed after Day 4 when acetate started to accumulate following ethanol fermentation (Figure 2: C). In the DSC group that received only ethanol, the relative abundance of *Methanothrix* within all archaeal taxa, calculated from sequencing data by summing the relative abundances of all *Methanothrix* ASVs, increased from 51 ± 0.4% on Day 4 to 55 ± 4% on Day 27 (Figure 2: C). This corresponded to approximately 18-fold increase in the absolute abundance of *Methanothrix* in the DSC bottles, from (9.7 ± 0.7) × 10^5^ copies/mL on Day 4 to (1.8 ± 0.2) × 10^7^ on Day 27 copies/mL (Figure 2: C). Other methanogens found in the GT2 culture were predominantly hydrogenotrophic, including *Methanoregula*, *Methanobacterium*, and *Methanospirillum* (Figure S5: B). Therefore, hydrogenotrophic methanogens in this study were approximated as all remaining archaeal ASVs other than acetoclastic *Methanothrix*. The decrease in the relative abundance of hydrogenotrophic methanogens in the EXP and DSC bottles could result from (1) the depletion of hydrogen due to reductive dechlorination and hydrogenotrophic methanogenesis; (2) accumulation of acetate from ethanol fermentation which facilitated acetoclastic methanogenesis; and (3) faster endogenous decay of microbial biomass as compared to acetoclastic methanogens.

### 3.2. Decay constant estimation from experimental data

In the experiment, the 16S rRNA gene copies of *Dhb* were directly quantified by qPCR, providing crucial information for estimating the decay constant for the dechlorinator. By assuming first-order decay kinetics during starvation, *Dhb* concentration data from nine bottles were used for decay constant estimation, where a clear trend of *Dhb* decay was observed. As shown in Table 2, the number of data points used for generating each estimate is *n*, with the standard error (SE), and adjusted R^2^ provided. For each group, the weighted mean and its combined standard error were derived by considering the individual SEs of each estimate within the group (Text S8). It is worth noting that the estimated *Dhb* decay constant (*b_Dhb_*) varied substantially across experimental groups, ranging from 0.024 day^-1^ in the EXP group to 0.088 day^-1^ in the DSC group. This variation might reflect the differences in growth conditions applied across groups. In particular, the faster decay of *Dhb* observed in the starved control groups (SC and DSC) may be attributed to intentional substrate limitation relative to the EXP group. This result highlights that the decay constant for a single microbial group within a mixed culture can vary with growth conditions, even though *b* is typically treated as a lumped parameter and assumed constant in biokinetics modelling. Decay dynamics in slow-growing anaerobic microorganisms can involve both endogenous and exogenous processes. Endogenous decay includes energy expenditures required to maintain essential cellular functions (e.g., transport, repair, and mobility) as well as loss of active cells due to aging.^68^ In contrast, exogenous decay reflects loss of cell activity caused by external factors such as viral predation, grazing by other microorganisms, or cell lysis under adverse environmental conditions (e.g., pH or temperature stress).^69^ The complexity of these processes likely contributes to the observed variability in decay behavior across different groups. In biokinetics modelling of enrichment cultures, such as GT2 in this study, decay is typically represented only through endogenous decay. In addition, changes in decay rate under starvation have been reported previously, where slower decay was observed during prolonged starvation (∼1000 days).^70^ This behaviour has been hypothesized to be linked to necromass recycling, which may partially offset biomass loss and reduce apparent decay over time.^70^

**Table 2.**
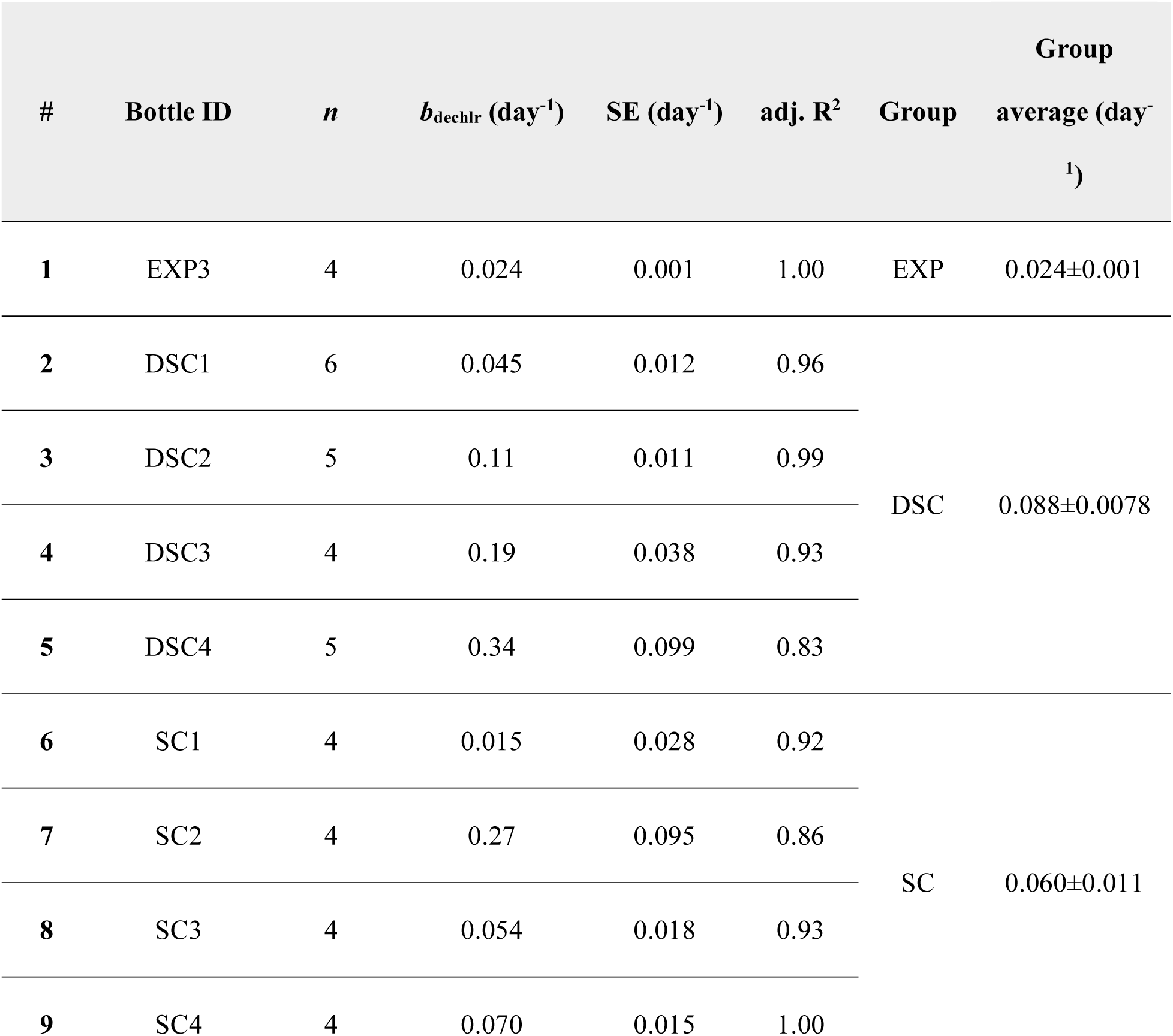
Decay constant estimations for *Dehalobacter* of the GT2 culture.

| # | Bottle ID | *n* | $b_{\text{dechl}r}$ (day<sup>-1</sup>) | SE (day<sup>-1</sup>) | adj. R<sup>2</sup> | Group | Group<br>average (day<sup>-1</sup>) |
| --- | --- | --- | --- | --- | --- | --- | --- |
| 1 | EXP3 | 4 | 0.024 | 0.001 | 1.00 | EXP | 0.024±0.001 |
| 2 | DSC1 | 6 | 0.045 | 0.012 | 0.96 | DSC | 0.088±0.0078 |
| 3 | DSC2 | 5 | 0.11 | 0.011 | 0.99 |  |  |
| 4 | DSC3 | 4 | 0.19 | 0.038 | 0.93 |  |  |
| 5 | DSC4 | 5 | 0.34 | 0.099 | 0.83 |  |  |
| 6 | SC1 | 4 | 0.015 | 0.028 | 0.92 | SC | 0.060±0.011 |
| 7 | SC2 | 4 | 0.27 | 0.095 | 0.86 |  |  |
| 8 | SC3 | 4 | 0.054 | 0.018 | 0.93 |  |  |
| 9 | SC4 | 4 | 0.070 | 0.015 | 1.00 |  |  |

Variations in the biomass of the key ethanol fermenter, Geobacterales_(o)_ASV1, were tracked by combining data collected from qPCR assays and 16S rRNA sequencing (Text S5). An arbitrary 10% relative uncertainty was assigned to relative abundance data obtained from 16S rRNA sequencing to account for instrumental and bioinformatic processing errors. Biomass data from bottles EXP4 and SC3 were used to estimate the decay constant of the ethanol fermenter (*b*_ferm_). Both decay constants estimated for these two groups were within the range for slow-growing heterotrophs (b < 0.07 day^-1^) at 25℃,^68^ supporting realistic estimation of biomass decay for ethanol fermenters in the model (Table 3).

**Table 3.** Decay constant estimations for the ethanol fermenter of the GT2 culture.

| # | Bottle ID | $n$ | $b_{\text{ferm}}$ (day<sup>-1</sup>) | SE (day<sup>-1</sup>) | adj. R<sup>2</sup> |
| --- | --- | --- | --- | --- | --- |
| 1 | EXP4 | 4 | 0.031 | 0.030 | 0.89 |
| 2 | SC3 | 5 | 0.045 | 0.050 | 0.77 |

Although bottles in the starved control (SC) group did not receive any external electron donor or acceptor after inoculation, where general biomass decay was expected, qPCR data revealed a non-decreasing trend in total bacterial and total archaeal gene concentrations, hindering the estimation of their respective decay constant. This phenomenon could result from (1) potential growth of hydrogenotrophic microorganisms supported by the hydrogen initially present in the bottle headspace; (2) endogenous decay of cells as roughly 80% of biomass were biodegradable^71^ and could be used by scavenger microorganisms for energy and growth; (3) the overestimation of active cell mass by qPCR, where undegraded genes from dead cells were picked up during the assay. For simplicity, neither factor was considered in the current model. After accounting for the temperature effect,^68^ a universal decay constant of 0.01 ± 0.01 day^-1^ at 25℃ was applied as the initial guess for all methanogens during parameter estimation, based on literature values for mesophiles and thermophiles in anaerobic digestors.^72–74^

### 3.3. Model simulation and parameter estimation

Using the parameter set that yielded the minimum WSSE in parameter estimation, model simulations showed good agreement with measured data (SSE ≤ 0.1, calculated using *r_i, j_* as defined in Section 2.7) for the total mass of lindane, MCB, and benzene, as well as for the dechlorinator biomass concentrations (Figure 3: A-C and E). A moderate fit (0.1 < SSE ≤ 0.5) was obtained for the biomass concentrations of ethanol fermenter (Figure 3: F). The fits of the remaining three model outputs used for parameter estimation, namely the total mass of methane, and biomass concentrations of acetoclastic and of hydrogenotrophic methanogens, were fair (SSE > 0.5). The model simulation failed to capture the growth phase within the first 30 days of the experiment of both types of methanogens as observed in measured data (Figure S8). The mismatches between simulated and measured data of model outputs related to methanogenesis were possibly caused by the presence of additional substrates for methanogenesis (e.g. acetate and hydrogen) in EXP2 that the model did not account for (Figure 3: D).

**Figure 3.**
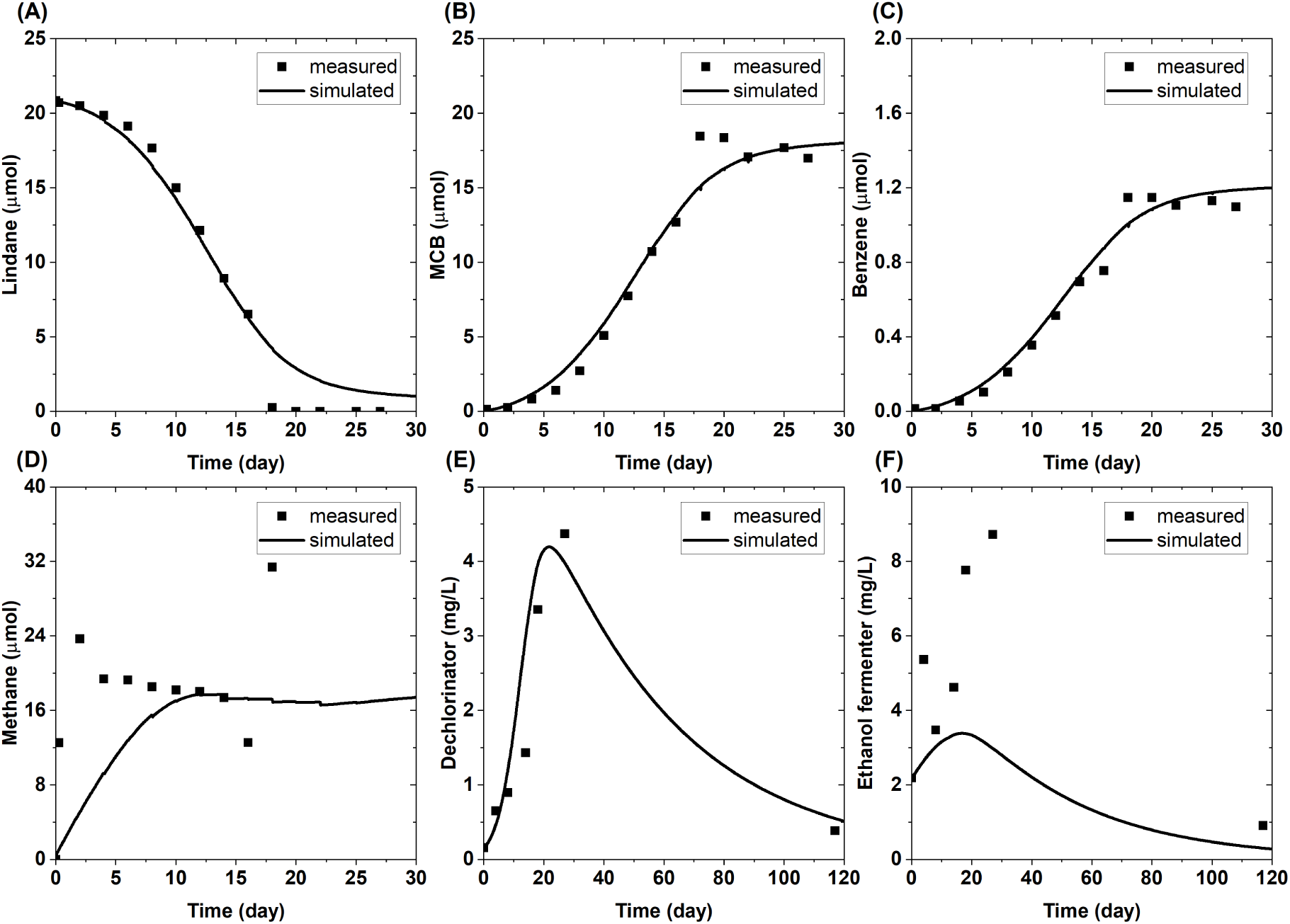
Measured and simulated chemical data and biomass data for bottle EXP2. A-D: total mass per bottle of lindane, MCB, benzene, and methane; E-F: biomass concentration of lindane dechlorinator and ethanol fermenter. Time range used for fitting for each data series correspond to those used in parameter estimation.

The parameter set that gives the lowest WSSE (i.e., the best fit) as well as the mean and standard deviation (SD) of each estimated parameter in all 101 BO runs are listed in Table 4. The imposed parameter bounds were based on previously reported values to restrict parameter estimation to physically and biologically reasonable ranges. The variability of the estimated parameters differed among parameters, as indicated by their coefficients of variation, calculated as the SD divided by the mean of the parameter estimates. For instance, this ratio is 22% for *k*_dechlr_, 50% for *b*_ferm_ and *b*_[meth, AC]_, and 99% for *K_s_*_[dechlr]_. For the estimated parameters, the greater their coefficients of variation, the less sensitive the model is towards them. The following section provides a more detailed and quantitative analysis of model sensitivity to these biokinetic parameters involved in this model.

**Table 4.**
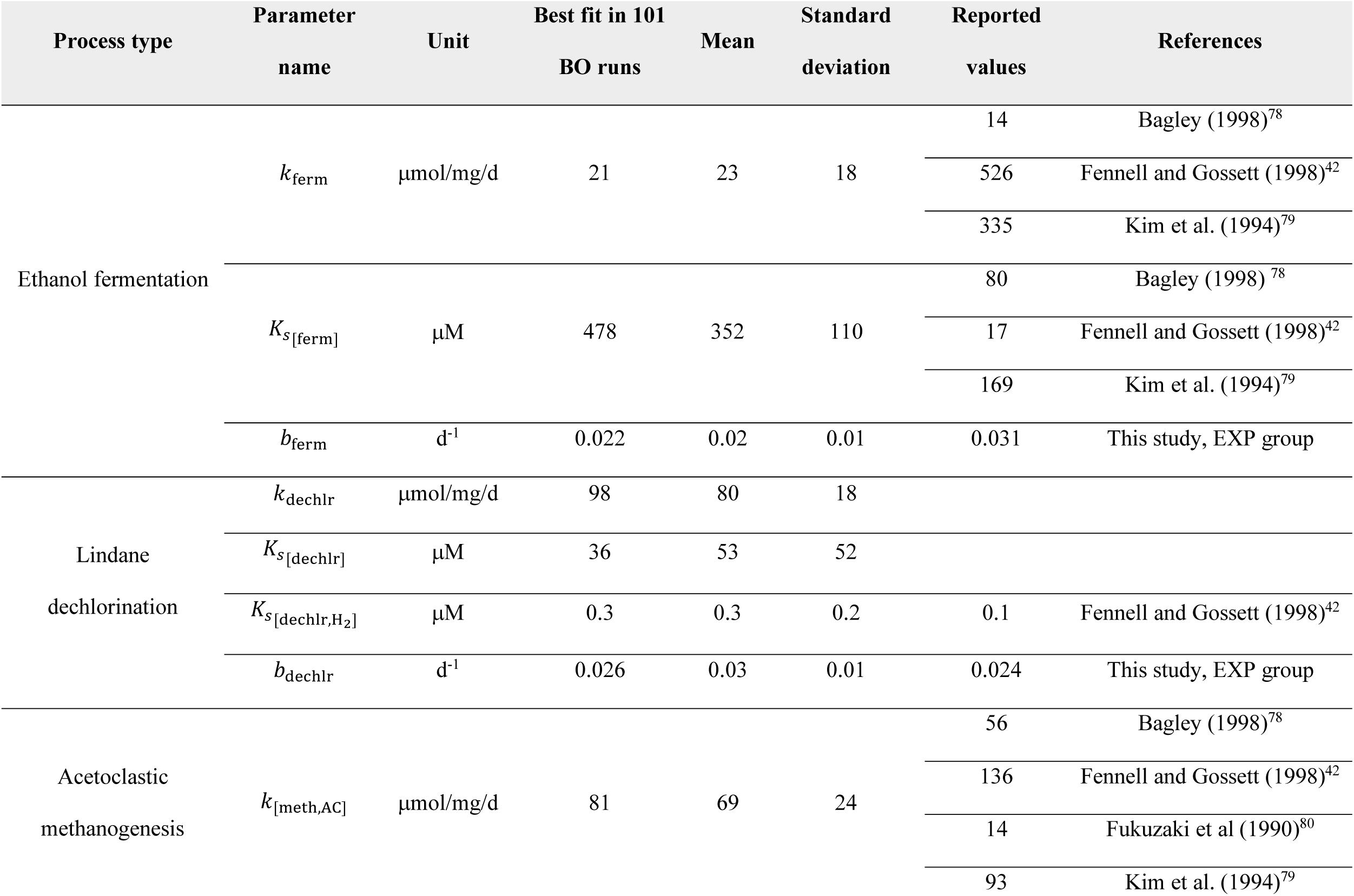
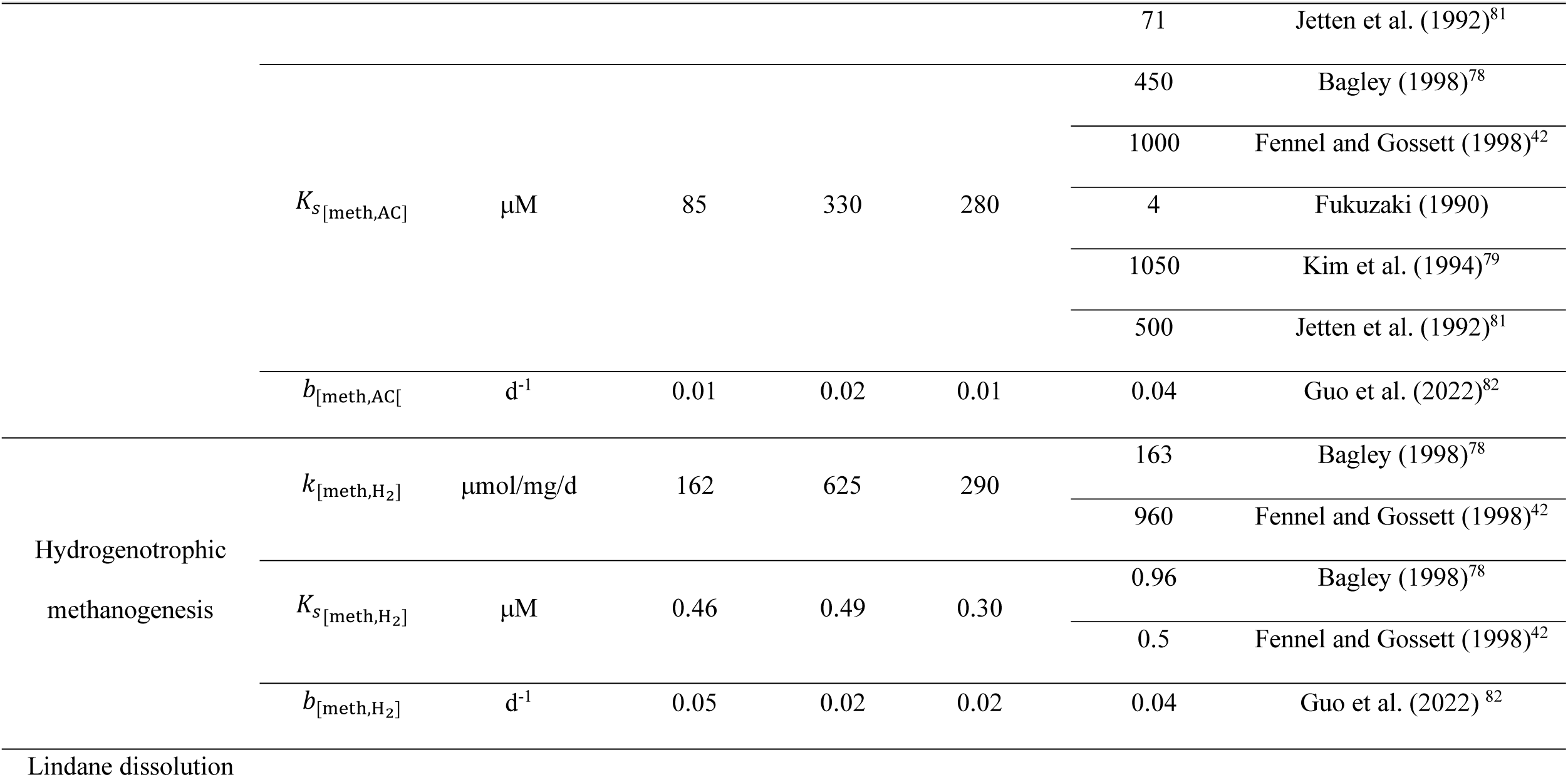
Parameter estimation results for the GT2 biokinetics model.

The range of biomass yields calculated through reaction thermodynamics as in the TEEM2 model (Text S9) is also compared against average biomass yields from previous biokinetics studies as shown in Table 5. The result shows comparable ranges between the two types of yields despite the maximum biomass yield output from the model being generally higher than previously reported average cell yields for the four microbial processes modelled. The minimum biomass yield was zero for acetoclastic and hydrogenotrophic methanogenesis in the simulation of the EXP2 scenario. A net zero biomass yield indicates limited energy generated from the process of substrate utilization (i.e., equals to or less than the Δ*G*_cr_) or the depletion of substrate for both processes. For the *Dhb* biomass yield during lindane dechlorination, model simulation outputs represent true yield as biomass decay is calculated separately. These modelled true yields were slightly larger than the net yields calculated from lab data of EXP2, which inherently incorporated cell decay. However, the simulated true yields were one to two orders of magnitude greater than previously reported values from other bench-scale experiments^12, 13^ (Table 5). This level of discrepancy between true yield and net yield is not uncommon, as rapid cell decay and sparse sampling frequency in lab studies may fail to capture the peak biomass concentrations during active microbial growth, leading to underestimation of biomass yields. Moreover, in bench-scale systems, the slow dissolution of lindane from the solid phase tends to maintain aqueous concentrations near the solubility limit of 9 mg/L (or 31 µM), supporting prolonged microbial growth before measurable declines in lindane concentration can be detected. The biokinetics model developed in this study considers both solid-phase dissolution and microbial dechlorination. The amount of lindane added in EXP2 far exceeded its solubility limit (∼20 µmol in 220 mL), providing a solid-phase reservoir that sustained lindane availability and prolonged the period of active dechlorination Because biomass yield was calculated dynamically throughout the simulation, this prolonged growth period contributed to higher average simulated yield. In contrast, the two prior bench-scale studies^12, 13^ that reported *Dhb* biomass yields amended lindane at concentrations closer to its solubility limit (∼3 - 4 µmol in 90 mL), likely providing more representative yield estimates when the lindane level is relatively low.

**Table 5.**
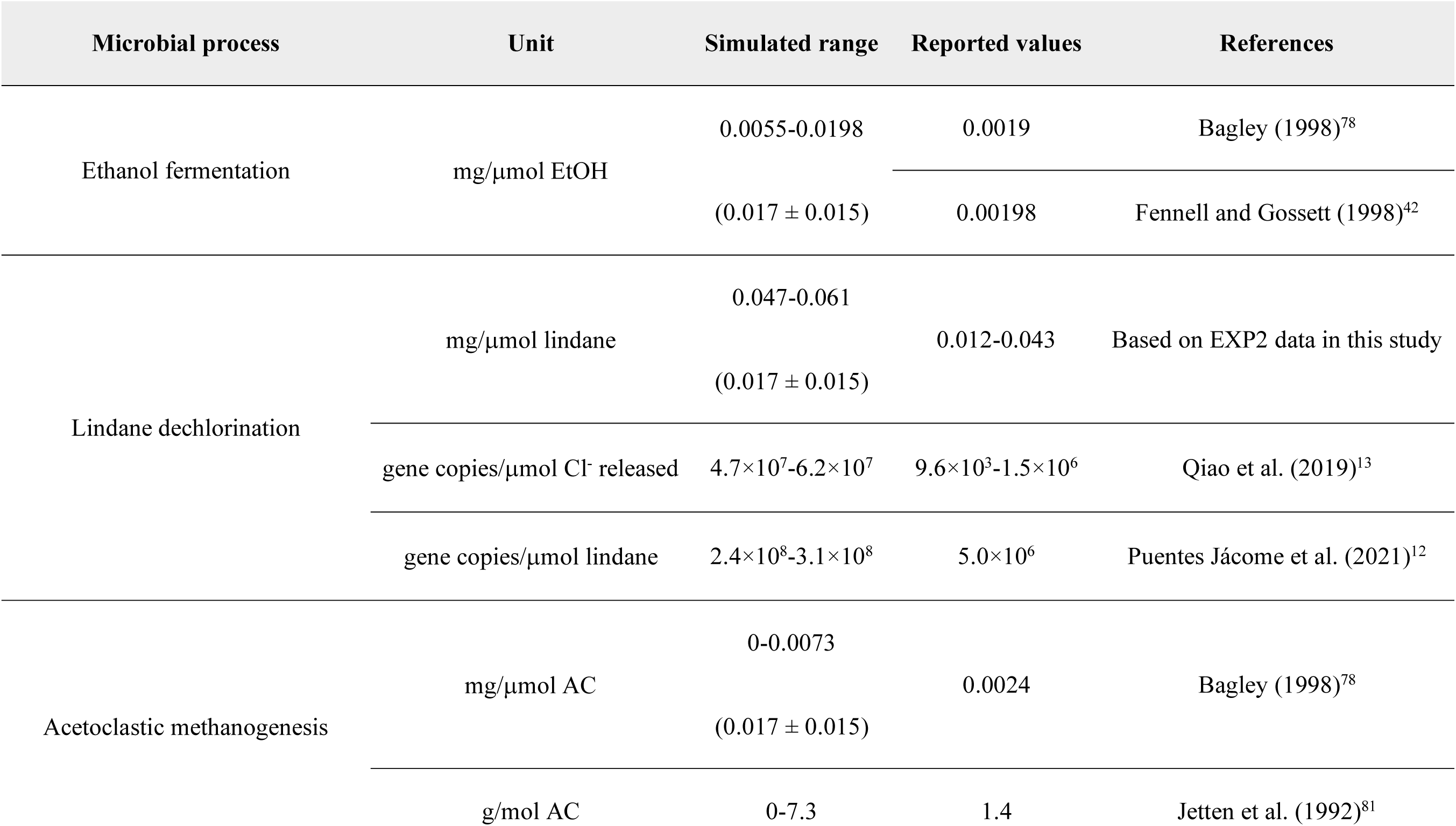
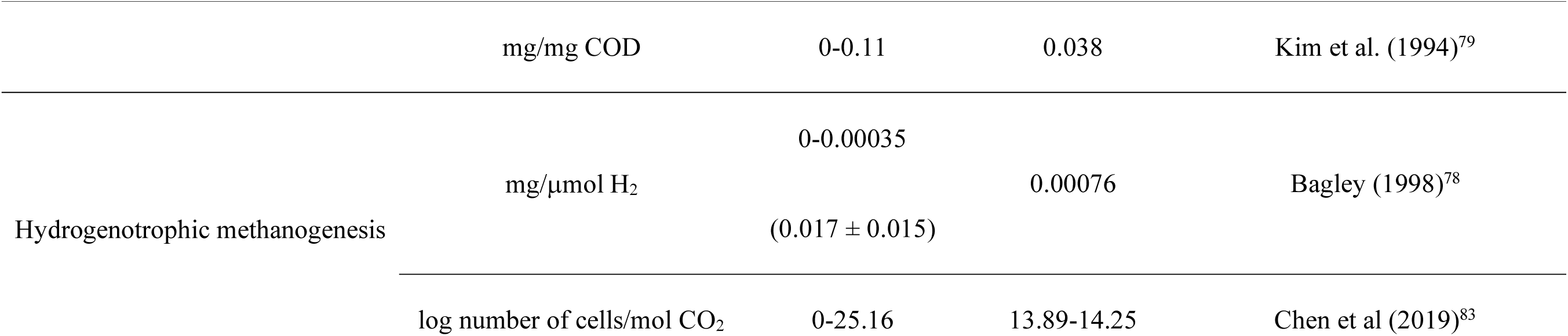
Range of cell yields for microbial processes in the GT2 biokinetics model. For the constant-yield scenario, values are presented as (mean ± standard deviation) of the cell yields estimated through parameter estimation. For both simulations, the parameter set of the lowest WSSE BO run was used.

In the primary model, biomass yields were calculated dynamically using the TEEM2 framework and therefore varied with reaction thermodynamics over the course of the simulation. For comparison, a constant-yield scenario was also evaluated, in which the biomass yield for each microbial group was fixed throughout the simulation and treated as an estimated parameter. These constant biomass yields were estimated using the same parameter estimation approach described in Section 2.7, with parameter ranges and initial guesses provided in Table S8. For each modelled microbial group except hydrogenotrophic methanogens, the estimated constant-yield ranges largely overlapped with the yields obtained from the variable-yield scenario corresponding to the simulation run with the lowest WSSE (Table 5). In contrast, the constant yield estimated for hydrogenotrophic methanogens was approximately one order of magnitude greater than the variable yield calculated using TEEM2. This suggests that yield estimates for hydrogenotrophic methanogenesis are more sensitive to TEEM2 assumptions than those for other microbial processes in this culture.

Based on carbon mass balance calculations (accounting for mass removal due to liquid sampling) of the EXP group, for the model input of ethanol, its maximum expected concentration should be 0.25 mM, which corresponds to a maximum acetate mass of 55 μmol/bottle due to fermentation. However, in all EXP bottles, their acetate concentration exceeded 100 μmol/bottle within the first day of the experiment, resulting in aforementioned higher than expected methane production (Figure 4: A-B). Clear inhibition of methanogenesis was also observed in the EXP group during the first 16 days compared to the DSC and SC groups, which could partially explain the concurrent buildup of acetate (Figure 4: A). Meanwhile, in the SC group, which did not receive any electron donors except for the initial hydrogen present in the anaerobic chamber (1%), its methane concentration kept increasing during the first 30 days of the experiment despite low acetate concentration (< 10 μmol/bottle or 0.03 mM, excluding SC1) (Figure 4: A-B). Methane and acetate concentrations from EXP and SC groups suggested (1) formation of acetate from processes besides ethanol fermentation when lindane was added as in the EXP group; (2) potential source of substrates for methanogenesis such as acetate (probably from biomass decay) and hydrogen (potentially from intrusion during sampling) as in the SC group, because both acetoclastic and hydrogenotrophic methanogens grew in the SC bottles. Moreover, for the DSC group, which only received ethanol after culture inoculation, its amount of methane production was greater than that of the EXP group and the SC group (Figure 4: A), even though its total acetate mass never exceeded 55 μmol/bottle, the expected amount based on carbon mass balance and ethanol feeding (Figure 4: B). VFAs measurements revealed the notable production of propionate only in the EXP group and the DSC group from around 0.22 μmol/bottle on Day 0 to up to 27 μmol/bottle on average by Day 4, yet the gradual consumption of propionate was more prominent in the DSC group. By Day 30, the average propionate mass in the EXP group was 20 μmol/bottle compared to 1.4 μmol/bottle for the DSC group (Figure 4: C). If preferential syntrophic propionate conversion to acetate and hydrogen happened in the DSC group,^75–77^ this discrepancy in propionate consumption between the EXP group and the DSC group could explain the relatively higher methane production in the DSC bottles.

**Figure 4.**
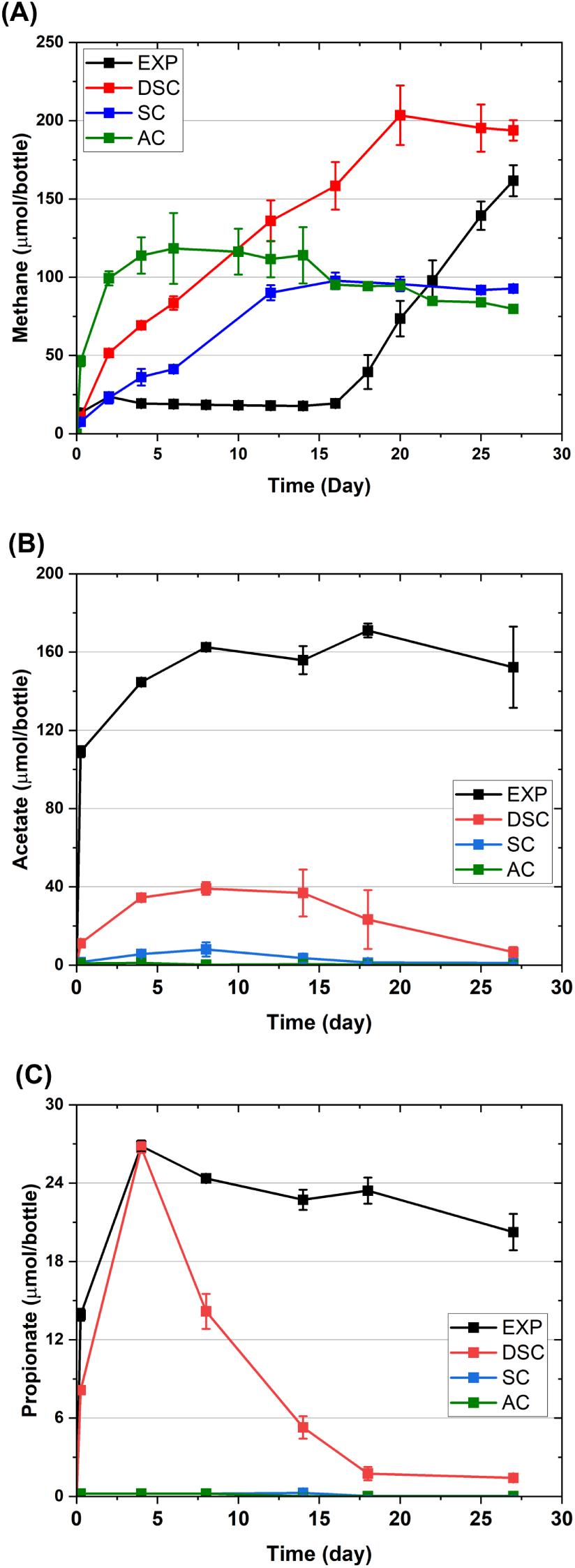
Masses of methane of acetate and propionate in the experimental (EXP), *Dehalobacter* starved control (DSC), starved control (SC), and analytical control (AC) groups. Error bars represent within-group standard deviations.

### 3.4. Sensitivity analysis results

The linear correlation between the global model output and values of individual parameters was quantified using standardized regression and partial correlation. Both methods flagged the same parameters as significant at the 0.05 level and in the same direction, serving as robust evidence for their unique impacts (Table 6). These influential parameters include *k* and *K_S_* for ethanol fermentation, lindane dechlorination, and acetoclastic methanogenesis. The *k* of hydrogenotrophic methanogenesis is also significant. This revealed the collective influence of parameters from all four modelled microbial processes and that no single process dominates the model’s behaviour. The fact that the partial correlation coefficients are larger in magnitude than the standardized regression coefficients can be due to the difference in each method when accounting for shared variance. The former measures the strength of the linear association between a parameter and the output after removing the contribution of all other parameters, while the latter estimates coefficients based on a full multiple linear regression model that can be affected by collinearity or overlapping variance. In terms of monotonic relationship between a parameter and the global output, Spearman correlation showed similar results as the approaches for linear relationship, but with some differences, as the *Ks* of ethanol fermentation (*K_S_*_[ferm]_) and of hydrogen during dechlorination (*K_S_*_[dechlr,H2]_) are no longer influential under the Spearman approach. The Morris screening method indicated significant (|EE| > 1) yet nonlinear response of the model to decay coefficients of all microbial processes as the microorganisms experienced both growth and decay in the simulation under the 120-day’s EXP2 scenario. Interestingly, *K_S_*_[dechlr,H2]_ was found to be quite significant under the Morris approach, signalling the nonlinearity of its impact on the model that cannot be fully captured by the Spearman approach.

**Table 6.**
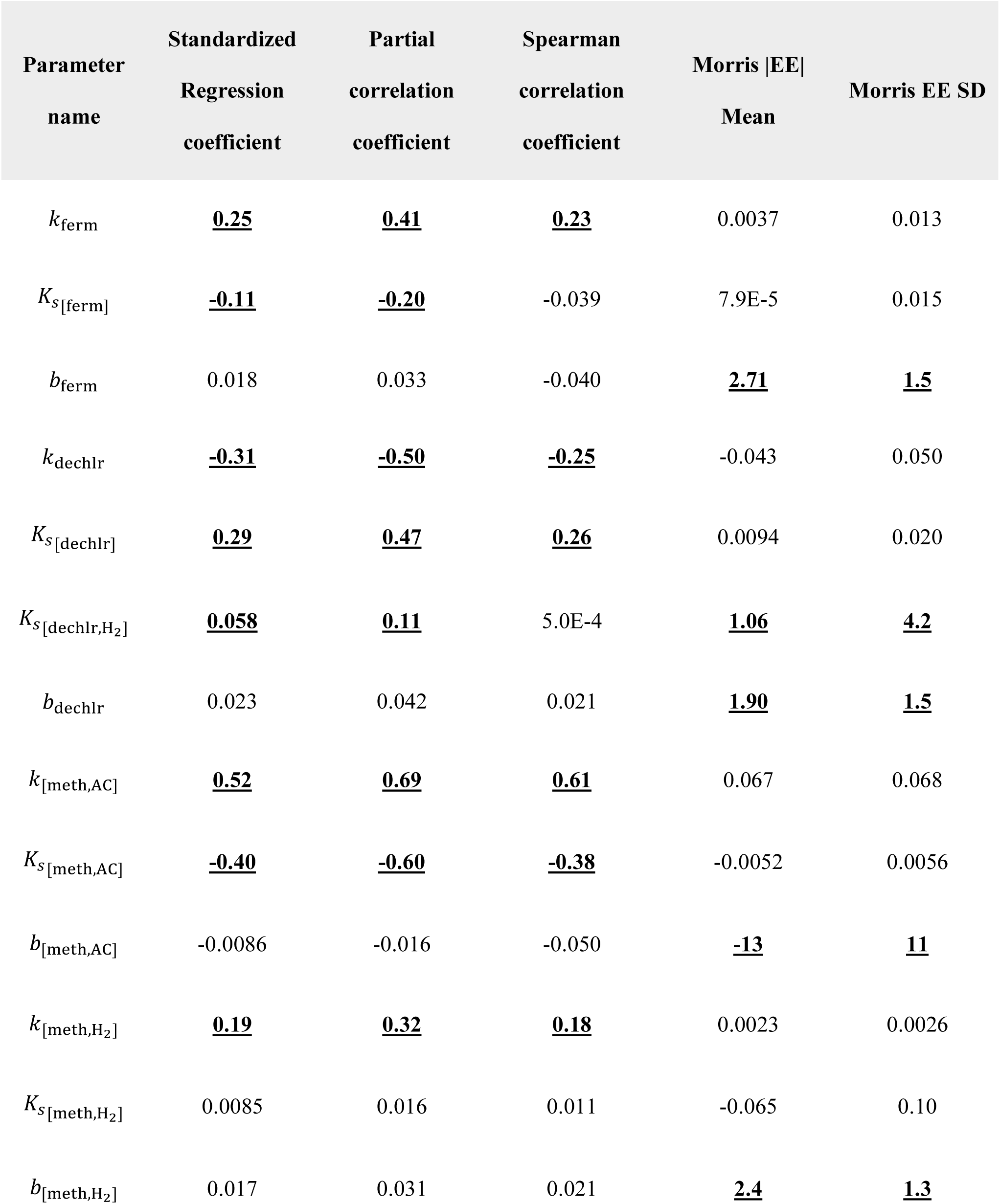
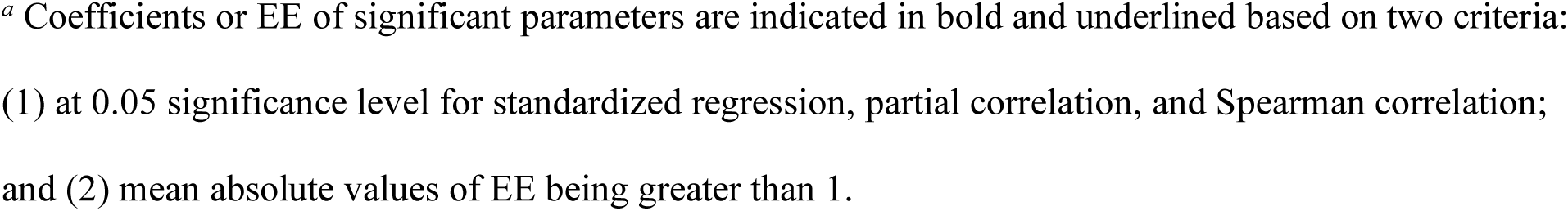
Global sensitivity analysis for response of global output to parameter values.*^as^*.

Evaluation of the responses of eight individual model outputs to varying parameter values using standardized regression (Figure S11), partial correlation (Figure S12), and Spearman correlation (Figure S13) consistently flagged the *k* and *K_s_* of the ethanol fermenter as significant. These two parameters influenced not only biomass of the ethanol fermenter itself, but also the biomass of *Dhb*, both types of methanogens, and the masses of lindane, MCB, and benzene. Meanwhile, all three sensitivity analysis approaches identified the *k* and *K_s_* associated with lindane dechlorination as significant in affecting the masses of lindane, MCB, and benzene, as well as the ethanol fermenter biomass. Moreover, the *k* of hydrogenotrophic methanogenesis was found to be a significant parameter, across all three analyses, impacting the masses of the reactant and products of dechlorination and the biomass of *Dhb*. Collectively, these results highlight the strong interdependence among the four key microbial processes in the GT2 culture, demonstrating how variation in one process can propagate and influence the outcomes of multiple others.

### 3.5. Model limitations and environmental implications

Because GT2 is a microbial consortium enriched from a contaminant field site, consisting of more than 30 different microbial groups involved in various biochemical processes, the four microbial processes modelled in this study represent only a subset of its total activity. For example, processes such as the additional release of acetate in the EXP group were not accounted for. This acetate production may have been related to induced cell decay linked to lindane toxicity, rather than hydrogenotrophic acetogenesis, given the low hydrogen levels in the culture. Additionally, the possible intrusion of hydrogen from the glovebox during sampling was not accounted for and the absence of hydrogen concentration measurements limited the ability to constrain hydrogen dynamics in the model. Inhibitory effects were not explicitly defined using inhibition models (e.g., Haldane kinetics), regardless of their possible types (e.g., self-inhibition, competitive, noncompetitive). Rather, they were assumed to be indirectly captured through thermodynamic constraints in the Gibbs free energy calculations based on real-time concentrations of reactants and products. However, this thermodynamic approach primarily characterizes competitive inhibition where multiple microbial processes (e.g., reductive dechlorination of lindane and hydrogenotrophic methanogenesis) compete for the same rate-limiting substrate, such as hydrogen. Meanwhile, model accuracy could be improved by incorporating variable activity coefficients that account for changes in ionic strength,^84^ as well as applying temperature and salinity corrections to Henry’s law constants.^85, 86^ Moreover, the first-order decay rate assumed to be constant for each microbial group in the GT2 model, may in fact vary with changes in growth conditions, as indicated by experimental results in Section 3.2. This suggests a need for future work to develop more detailed mathematical representations of microbial decay in biokinetics models, potentially linking decay behaviour to parameters that describe growth conditions. By deriving kinetic parameters through nonlinear regression of experimental data for lindane dissolution and biomass decay, along with estimating Monod kinetics parameters for key microbial processes using Bayesian optimization, the GT2 biokinetics model effectively simulates the dissolution and anaerobic biotransformation of lindane. Despite the limitations discussed above, the model successfully captured the dynamics of syntrophic microbial growth in GT2, underscoring the critical inter-relationships among the four major functional microbial groups in determining degradation outcomes. By providing the first complete set of biokinetics parameters for the key microbial processes involved (Table 4), this study helps bridge the gap between batch-scale culture research and field-scale applications of anaerobic lindane biodegradation. Further study on culture behavior, especially lindane toxicity and mechanisms for potential growth inhibition, can improve model accuracy, refine simulations and boost model reliability in diverse environments. With the advancement of bioremediation strategies targeting degradation of MCB and benzene, the sequential anaerobic transformation of lindane into non-toxic end products is now a demonstrated possibility.^12^ Results from this study offer valuable insights for future research aimed at predicting and optimizing bioremediation performance under more complex and variable subsurface field conditions. For instance, with information on groundwater lindane concentration and microbial abundances during treatment, the model can be used to estimate remediation timelines, calculate electron donor dosage, and predict methane production from bioremediation.

