## Supplementary Information 1 for "Thermodynamically controlled biokinetics modelling of a lindane-degrading anaerobic enrichment culture"

### Supplementary Information for: Thermodynamically controlled biokinetics modelling of a lindane-degrading anaerobic enrichment culture

**Jingzhi Liang,[1,2] Line Lomheim,[2] Luz, A. Puentes Jácome, [3] Elizabeth A. Edwards,[2] Brent E. Sleep[1*]**

[1]Department of Civil & Mineral Engineering, University of Toronto, 35 Saint George Street, Toronto, Ontario, M5S 1A4, Canada

[2]Department of Chemical Engineering and Applied Chemistry, University of Toronto, 200 College Street, Toronto, Ontario, M5S 3E5, Canada

[3]Department of Biology, York University, 115 Ottawa Road, North York, Ontario, M3J 1P3, Canada

**Corresponding Author:**

Number of Pages: 37

Number of Supporting Texts: 9

Number of Tables: 16

Number of Figures: 16

#### List of Supplementary Texts

Figure S6. Microbial community structure at the amplicon sequence variants (ASV)level for bottles EXP2, EXP3 and EXP4 of the experimental (EXP) group: (A) Bacteria, (B) Archaea. .. 25

Figure S12. Partial correlation analysis of individual model output, including mass of lindane ($m_{lin}$), MCB ($m_{MCB}$), benzene ($m_{BZ}$), methane ($m_{CH4}$), and biomass concentrations of *Dehalobacter* ($X_{Dhb}$), ethanol fermenter ($X_{ferm}$), acetoclastic methanogens ($X_{AC\_meth}$), and hydrogenotrophic methanogens ($X_{H2\_meth}$). Asterisks mark parameters that are statistically significant at the 0.05 level. ........................................................................................ 31

Figure S13. Spearman correlation analysis of individual model output, including mass of lindane ($m_{lin}$), MCB ($m_{MCB}$), benzene ($m_{BZ}$), methane ($m_{CH4}$), and biomass concentrations of *Dehalobacter* ($X_{Dhb}$), ethanol fermenter ($X_{ferm}$), acetoclastic methanogens ($X_{AC\_meth}$), and hydrogenotrophic methanogens ($X_{H2\_meth}$). Asterisks mark parameters that are statistically significant at the 0.05 level. ........................................................................................ 32

Figure S14. Example Simulink model screenshot: thermodynamics block for lindane dechlorination. Signal flow follows the direction of the connected arrows. Regular and rounded rectangular symbols denote input or output scalar signals, symbols with the MATLAB icon denote coding blocks, and scope symbols denote signal-monitoring blocks for tracking output values. ........................................................................................ 33

Figure S15. Example Simulink model screenshot: biokinetics block for lindane dechlorination. Signal flow follows the direction of the connected arrows. Regular and rounded rectangular symbols denote input or output scalar signals, symbols with the MATLAB icon denote coding blocks, scope symbols denote signal-monitoring blocks for tracking output values, and symbols with integrator signs denote integration of input signals. ........................................................ 33

Figure S16. Example Simulink model screenshot: mass balance block for MCB and benzene. Signal flow follows the direction of the connected arrows. Regular and rounded rectangular symbols denote input or output scalar signals, symbols with the MATLAB icon denote coding blocks, scope symbols denote signal-monitoring blocks for tracking output values, symbols with integrator signs denote integration of input signals, and symbols with multiplication signs denote multiplication of input signals. ........................................................................................ 34

#### Text S1. Lindane measurements

Lindane concentration was determined using a gas chromatography with mass spectrometry (GC-MS) system (Agilent 7890B and 5977B GC/MSD) equipped with a DB-5ms column (dimensions: 30 m × 250 µm × 0.25 µm) and an Agilent PAL RSI 85 autosampler. Samples were extracted using Solid Phase Micro Extraction (SPME) with a 50/30 µm DVB/CAR/PDMS fiber (Supelco, Sigma-Aldrich), where a 10 mL diluted sample (0.5 mL liquid sample dissolved in 9.5 mL acidified 10 w/v% NaCl solution, pH = 2.5) was exposed to the fiber for 28 minutes at 60°C while being agitated at 250 rpm. The GC injection port was set to 250°C in splitless mode with helium as the carrier gas flowing at a rate of 1.0 mL/min. The oven temperature program started at 70°C, ramped to 210°C at 10°C/min (held for 1 minute), and then ramped to 285°C at 25°C/min (held for 1 minute). The MS operated in the electron ionization positive mode (EI+) with an ion source temperature of 230°C, quadrupole temperature of 150°C, transfer line temperature of 250°C, electron multiplier voltage of 1164 V, electron energy 70 eV, and a mass spectra m/z scan range between 40 and 350. Calibration involved preparing aqueous lindane standards (0.1 mg/L – 10 mg/L), and quantification was performed by comparing the peak areas in samples to a calibration curve. Quality control included method blanks, and data analysis was conducted using Agilent ChemStation. All lindane data measured are presented in Figure S2, with raw data reported in Table S12.

#### Text S2. Volatile organic compounds measurement

Monochlorobenzene, benzene and methane were quantified using a gas chromatograph equipped with a flame ionization detector (GC-FID). During sampling, liquid culture samples (0.5 mL or 1 mL) were mixed with 5 mL of MilliQ water acidified to a pH below 2 using 6N HCl to inhibit microbial activity. Samples were then equilibrated in an Agilent G1888 headspace autosampler at 70°C for 40 minutes. Following equilibration, 3 mL headspace samples were automatically injected into an Agilent 7890A GC equipped with an Agilent GS-Q column (30 m length, 0.53 mm inner diameter). Helium served as carrier gas at a flow rate of 11 mL/min. The injector and detector temperatures were set to 200°C and 250°C, respectively. The oven temperature program started at 35°C for 1.5 minutes, increased to 100°C at 15°C/min, then to 185°C at 5°C/min (held for 10 minutes), and then followed by a ramp to 200°C at 20°C/min (held for 10 minutes). MCB and benzene data from the experimental (EXP) group and the analytical control (AC) group are presented in Figure S3, with raw data reported in Table S12.

**Text S3. Anion measurements**

The method for measuring anions of volatile fatty acids acetate and propionate using a Dionex Integrion High-Pressure Ion Chromatography (HPIC) system (Thermo Scientific) involved the use of a Dionex AS-DV autosampler, a Dionex IonPac AS11-HC Analytical Column (2× 250 mm) and a Dionex IonPac AG11-HC Guard Column (2 × 50 mm). The eluent was a potassium hydroxide (KOH) solution (Dionex EGC 500 KOH eluent generator cartridge) with a multi-step gradient that set off at 0.50 mM, ramped up to 2.50 mM KOH at 0.20 mM/min, then increased to 30 mM at 1.4 mM/min (held for 10 minutes), then ramped down to 0.50 mM at 15 mM/min (held for 5 minutes). Anion standards were prepared in MilliQ water, ranging from 0.001 mM to 0.5 mM for acetate and propionate. During sample preparation, samples were filtered using 0.2 μm filters, diluted 10 times, and prepared in 0.5 mL PolyVials with filter caps (Thermo Scientific). The column temperature is set to 30°C, with a pump rate of 0.25 mL/min and an injection volume of 500 μL. Detection was done using suppressed conductivity mode with the suppressor current automatically controlled by the instrument. Data processing was performed using the Chromeleon 7 Chromatography Data System software. Calibration standards and blank samples were run at regular intervals to ensure accuracy and to check for contamination or carryover.

**Text S4. DNA extraction and quantitative PCR**

DNA extractions were carried out using a MagMAX Microbiome Total Nucleic Acid Kit (Thermo Fisher Scientific) and a DNA extraction robot: KingFisher Duo Prime (Thermo Fisher Scientific) following manufacturer's protocol. For each DNA sample, 2 mL of culture liquid was harvested. The liquid sample was then centrifuged for 20 mins at 13,000 g and 10 °C to collect the cell pellet with supernatant removed. The cell pellet was subsequently stored at -80°C prior to DNA extraction. Purified DNA samples were eluted into the elution buffer provided in the DNA extraction kit at 80 μL/sample. Extraction efficiency was assessed through DNA concentration measurements using an Invitrogen Qubit 3.0 Fluorometer (Thermo Fisher Scientific) and an Invitrogen Qubit dsDNA High-Sensitivity Assay Kit. The minimum quantifiable DNA concentration in a sample elution is 0.005 ng/μL. For each purified DNA sample, quantitative polymerase chain reaction (qPCR) assays were carried out to determine the absolute concentrations of total bacterial genes (GenBac), total archaeal genes (GenArch) and *Dehalobacter* genes (*Dhb*) within using their respective 16S rRNA gene primers (Table A1). qPCR assays were prepared in duplicates inside of a PCR UV cabinet (Esco Technologies). A series of IDTE buffer (Integrated DNA Technologies)-diluted plasmid DNA with known gene concentrations ranging from 10 to $10^8$ gene copies/µL ($10^6$ gene copies/µL for *Dhb*) were also prepared as calibration standards for every qPCR run (Figure S2). For each qPCR reaction, 2 µL DNA template, 1 µL forward primer (10 mM), 1 µL reverse primer (10 mM), 10 µL SsoFast EvaGreen Supermix (Bio-Rad Laboratories), and 6 µL UV-sterile UltraPure DNase/RNase-Free Distilled Water (Thermo Fisher Scientific) were used. qPCR reactions were analyzed on a C1000 Thermal cycler coupled with a CFX96 real-time PCR detection system (Bio-Rad Laboratories), and the Bio-Rad CFX Manager (v 3.1) software was used for initial data processing. The thermal cycling programs for each qPCR target are provided in Table S2. Calibration information for qPCR assays is presented in Figure A1. The amplification efficiencies for all qPCR runs ranged from: 86.6%-98.4% for *Dhb*, 98.5%-103.1% for GenBac, and 93.3%-100.2% for GenArch.

**Text S5. Biomass concentration calculations**

In this study, data from next-generation sequencing (NGS) for 16S rRNA genes from all microorganisms were used to derive the relative abundances of individual microbial groups within all bacterial or archaeal taxa. Notably, considering potential differences in the sensitivity of bacterial and archaeal 16S rRNA genes to the primer set used in NGS, the elative abundances of bacterial and archaeal taxa were calculated separately for the two domains. Meanwhile, qPCR assays from DNA samples provided concentrations of total bacterial 16S rRNA genes and total archaeal 16S rRNA genes. By combining these datasets, the biomass (in 16S rRNA gene copies/mL) of individual groups of bacteria or archaea identified in NGS can be calculated as follows:

$$\text{Bacterial biomass concentration } \left({}^{\text{gene copies}}/_{\text{mL}}\right) =$$

$$\text{Relative abundance in all bacterial taxa (\%)} \times \text{total bacterial gene concentration } \left({}^{\text{gene copies}}/_{\text{mL}}\right)$$

Or

Archeal biomass concentration $\left({}^{\text{gene copies}}/_{\text{mL}}\right)$ = Relative abundance in all archaeal taxa (%) × total archaeal gene concentration $({}^{\text{gene copies}}/_{\text{mL}})$

**Text S6. Dry cell biomass calculations**

The general cell shape and dimensions and the number of 16S rRNA gene/cell vary between different microbial groups (Table S3). Therefore, additional calculations are required to convert biomass concentrations in *gene copies/mL* to dry cell biomass in *mg dry mass/mL* using the representative microorganism in each microbial group modelled in this study. The conversion from gene copy number to dry cell biomass is carried out assuming an average wet cell density[1] of 1.1571 g/mL and 40% of the wet cell mass is dry mass[2].

**Text S7. Lumped mass transfer rate coefficient ($K_l$) estimation**

Aqueous lindane concentrations measured in the three AC bottles were used to acquire the initial guess of $K_l$ for lindane. Parameter estimation was carried out using the Nonlinear Curve Fit tool in OriginPro (2023b). The first-order dissolution kinetics is described as follows:

$$C_{lin} = C_{s[lin]} \cdot (1 - e^{-K_l \cdot t})$$

where $C_{lin}$ denotes transient lindane concentration, $C_{s[lin]}$ is the solubility limit of lindane under experimental conditions, which was set as maximum lindane concentration measured in the AC bottles (9653.24 µg/L) for parameter initialization. During the estimation process, the average $C_{lin}$ of all AC bottles between Day 0 and Day 14 was weighted by the standard deviation (SD) of $C_{lin}$ as $\frac{1}{SD^2}$. The fit converged when $\chi^2$ tolerance reached $10^{-9}$ in 17 iterations, with an adjusted $R^2$ of 0.67. The resultant estimated parameters are $K_l = 2.4 \pm 0.96$ day$^{-1}$, $C_{s[lin]} = 9.0 \pm 0.16$ mg/L (Figure S1).

**Text S8. Decay constant estimation from experimental data**

Nonlinear curve fitting was performed assuming simple first-order decay of target microbial groups during starvation. The decay function is expressed as:

$$y = y_0 + Ae^{-bx}$$

where $y$ is the biomass concentration at time $x$ (in days), $y_0$ is the minimum biomass concentration, $A$ is the maximum biomass concentration (or the initial value of the decay process), and $b$ is the decay constant in day$^{-1}$.

In each nonlinear curve fitting run, three parameters were estimated, with initial values set as $y_0$ = 0 gene copies/mL, A = maximum biomass concentration acquired using molecular biology tools, and $b$ = 0.1 day$^{-1}$. Each input concentration data ($y_i$) was weighted by its error ($e_i$) as $\frac{1}{e_i^2}$. Levenberg-Marquardt algorithm was used to determine the best-fitted parameter set, achieving a $\chi^2$ tolerance value of $10^{-9}$ within 400 iterations. The standard error for each estimate ($\sigma_i$) was scaled by the squared root of reduced $\chi^2$.

To calculate an average decay constant ($\bar{b}$) from multiple estimates ($b_i$) within a group, the weighted mean of all estimates was computed as:

$$\bar{b} = \frac{\Sigma(\frac{b_i}{\sigma_i^2})}{\Sigma(\frac{1}{\sigma_i^2})}$$

while the combined error was calculated to account for the standard error of each estimate as:

$$\sqrt{1/\Sigma(\frac{1}{\sigma_i^2})}.$$

For the estimation of the *Dhb* decay constant ($b_{Dhb}$), concentration errors $e_i$ were equal to measurement errors from *Dhb* qPCR assays. For the estimation of the decay constant for the key ethanol fermenter Geobacterales_(o)_ASV1 ($b_{ferm}$), a different approach was used to acquire concentration errors. This was because the concentration of this fermenter was calculated as the product of GenBac qPCR and relative abundance in all bacterial taxa. For simplicity, an arbitrary 10% relative error ($r_a$) was assigned to relative abundances derived from sequencing data. And the relative error of GenBac qPCR ($r_{qPCR}$) can be calculated as the ratio between the standard deviation and mean in duplicated qPCR assays for each DNA sample. To account for error propagation due to multiplication, the relative error of each concentration data point ($r_i$) was calculated as

$$r_i = \sqrt{r_a^2 + r_{qPCR}^2}$$

and $e_i$ was determined as the product of $y_i$ and $r_i$.

**Text S9. Cell yield calculations**

Yields of the four modelled microbial processes were calculated during simulation based on Gibbs free energies of both metabolic and anabolic reactions. Using the McCarty efficiency approach,[3-5] the overall reaction ($R$) for each microbial process can be expressed as:

$$R = f_e R_a + f_s R_c - R_d$$

where $R_a$ is the electron acceptor half-reaction, $R_c$ is the cell synthesis half reaction, and $R_d$ is the electron donor half reaction. Here, $f_e$ is the fraction of electrons transferred from the electron donor to the electron acceptor for energy generation, while $f_s$ is the other fraction diverted to cell synthesis. By definition, $f_e + f_s = 1$.

When converted into mass units, such as *mg dry mass/μmol substrate*, $f_s$ is also termed the net yield ($Y_n$). $Y_n$ can be directly measured experimentally and is smaller than true yield ($Y$) due to cell decay:

$$Y_n = \frac{dX_a/dt}{-dS/dt} = Y - b \cdot \frac{X_a}{(-\,dS/dt)}$$

where $dX_a/dt$ is the net growth rate of active cell mass, $(-\,dS/dt)$ represents the substrate utilization rate, and $b$ is the decay constant for cells. $f_s^0$ is the unitless form of $Y$ and can by calculated through reaction energetics as follows:

$$f_s^0 = (1 - \frac{\frac{\Delta G_{ic}}{\varepsilon^n} + \frac{\Delta G_{pc}}{\varepsilon}}{\varepsilon \Delta G_e})^{-1}$$

According to the TEEM2 model,[5] $\Delta G_{ic}$ is the Gibbs free energy associated with converting the carbon source to acetyl-CoA, the intermediate compound for cell synthesis. The superscript $n$ relates to the sign of $\Delta G_{ic}$. If $\Delta G_{ic} > 0$, then $n = 1$, meaning energy is required to convert the carbon source to acetyl-CoA; otherwise, $n$ = -1. Based on Gibbs free energy of half-reactions as indicated in Table B6, under standard conditions, for the two carbon sources utilized by modelled microbial groups, ethanol and acetate, $\Delta G_{ic}$ equals 0.55 kJ/ $e^-$eq and 4.10 kJ/ $e^-$eq, respectively. $\varepsilon$ is the energy transfer efficiency during cell synthesis, which was set equal to 0.6 in the model.[4] $\Delta G_{pc}$ is the energy required for cell synthesis. Assuming a universal cell composition of $C_5H_7O_2N$, $\Delta G_{pc}$ is estimated to be 18.8 kJ/$e^-$eq when ammonia served as the nitrogen source.[4] $\Delta G_e$ is the instantaneous Gibbs free energy released from the energy reaction, which varies with the concentrations of reactants and products. The true fraction of electrons transferred for energy production $f_e^0 = 1 - f_s^0$.

True yield calculations in the model expanded the McCarty efficiency approach[3-5] by incorporating the general degree of reductance (GDR) of specific reaction participants. For a generic organic compound $C_nH_aO_bN_cP_dS_eCl_f$, its GDR = 4n+a-2b-3c+5d-2e-f. The calculation approach considered two scenarios related to the four microbial processes modelled:

(1) Single substrate ($S$) acting as both the carbon source and the energy source (e.g., acetoclastic methanogenesis and ethanol fermentation):

$$Y(\frac{mg\ dry\ cell\ mass}{\mu mol\ S}) = f_s^0 \cdot \frac{GDR_S}{GDR_C} \cdot \frac{M_{cell}}{10^3}$$

(2) Single substance acting as the electron donor ($D$, e.g, $H_2$) with a different compound (e.g, acetate) utilized as the carbon source (e.g., lindane dechlorination and hydrogenotrophic methanogenesis):

$$Y(\frac{mg\ dry\ cell\ mass}{\mu mol\ D}) = \frac{f_s^0}{f_e^0} \cdot \frac{GDR_D}{GDR_C} \cdot \frac{M_{cell}}{10^3}$$

In the two equations above, $M_{cell}$ is the molar mass of cells, which is 113.13 g/mol for the generalized cell formula $C_5H_7O_2N$, and GDR of cells ($GDR_C$) is 20.

**Table S1. Treatment table for the biokinetics experiment.**

| Group | Bottle ID | Treatment | | |
|---|---|---|---|---|
| | | 20% inoculation | Lindane | Ethanol |
| Experimental (EXP) | GT2-LIN_EtOH_ino_1 (EXP1) | x | x | x |
| | GT2-LIN_EtOH_ino_2 (EXP2) | x | x | x |
| | GT2-LIN_EtOH_ino_3 (EXP3) | x | x | x |
| | GT2-LIN_EtOH_ino_4 (EXP4) | x | x | x |
| Starved Control (SC) | GT2-ino_1 (SC1) | x | | |
| | GT2-ino_2 (SC2) | x | | |
| | GT2-ino_3 (SC3) | x | | |
| | GT2-ino_4 (SC4) | x | | |
| *Dhb* Starved Control (DSC) | GT2-EtOH_ino_1 (DSC1) | x | | x |
| | GT2-EtOH_ino_2 (DSC2) | x | | x |
| | GT2-EtOH_ino_3 (DSC3) | x | | x |
| | GT2-EtOH_ino_4 (DSC4) | x | | x |
| Analytical Control (AC) | GT2-LIN_EtOH_1 (AC1) | | x | x |
| | GT2-LIN_EtOH_2 (AC2) | | x | x |
| | GT2-LIN_EtOH_3 (AC3) | | x | x |

**Table S2. Quantitative PCR primer sequences.**

| Target microbial group | Primer set ID | Primer Sequence 5'-3' | Reference |
|---|---|---|---|
| *Dehalobacter* | Dhb 477f | GATTGACGGTACCTAACGAGG | Grostern and Edwards (2006)[6] |
| | Dhb 647r | TACAGTTTCCAATGCTTTACG | |
| General Bacteria | Bac_1055f | ATGGCTGTCGTCAGCT | Ferris et al. (1996),[7] Amann et al. (1995)[8] |
| | Bac_1392r | ACGGGCGGTGTGTAC | |
| General Archaea | Arch_787f | ATTAGATACCCGBGTAGTCC | Yu et al. (2005)[9] |
| | Arch_1059r | GCCATGCACCWCCTCT | |

#### Table S3. Thermal cycling programs for quantitative PCR assays.

| Target microbial group | Annealing temperature (°C) | Thermal cycling program |
|---|---|---|
| *Dehalobacter* | 62.5 | Denaturation at 98°C for 2 minutes, followed by 39 cycles of 5 sec at 98°C and 10 sec at 62.5°C (ramp rate of 2°C/sec). |
| General Bacteria | 55 | Denaturation at 98°C for 2 minutes, followed by 39 cycles of 5 sec at 98°C and 10 sec at 55°C (ramp rate of 2°C/sec). |
| General Archaea | 60 | Denaturation at 98°C for 2 minutes, followed by 39 cycles of 5 sec at 98°C and 10 sec at 60°C (ramp rate of 2°C/sec). |

#### Table S4. Cell information of modelled microbial groups for biomass conversion calculations.

| Microbial Group | Representative organism | Shape | Cell length (m) | Cell radius (m) | Cell Volume ($m^3$) | 16S gene copies/cell | Dry mass/gene copy (mg/copy) | References |
|---|---|---|---|---|---|---|---|---|
| Dechlorinator | *Dehalobacter* | cylinder | 2.50E-06 | 4.00E-07 | 1.26E-18 | 3 | 1.94E-10 | Holliger et al. (1998),[10] Jacome and Edwards (2017)[11] |
| Ethanol fermenter | *Geobacter* | cylinder | 2.00E-06 | 5.00E-07 | 1.57E-18 | 3 | 2.42E-10 | Bansal et al. (2015),[12] Jacome and Edwards (2017)[11] |
| Hydrogenotrophic methanogen | *Methanoregula* | cylinder or coccoid | 2.00E-06 | 2.50E-07 | 3.93E-19 | 1 | 1.82E-10 | Bauer et al. (2011),[13] Saengkerdsub et al. (2007)[14] |
| Acetoclastic methanogen | *Methanothrix* | cylinder | 6.25E-06 | 2.50E-07 | 1.23E-18 | 1 | 5.68E-10 | Mori et al. (2012),[15] Saengkerdsub et al. (2007)[14] |

**Table S5. Gibbs free energies of formation of substances related to thermodynamic calculations in this study.**

| Substance | State | $\Delta G_f^{\circ}$ (25°C), kJ/mol | Reference |
|---|---|---|---|
| $H_2$ | g | 0 | Thauer et al. (1997)[16] |
| $H^+$ (pH7) | aq | -39.87 | |
| $H_2O$ | liq | -237.178 | |
| $CO_2$ | aq | -386.02 | |
| $CO_2$ | g | -394.359 | |
| $HCO_3^-$ | aq | -586.85 | |
| $CH_4$ | g | -50.79 | |
| Ethanol | aq | -181.75 | |
| Acetate | aq | -369.41 | |
| Benzene | liq | 129.7 | Perry et al. (1984)[17] |
| | aq | 133.9 | Rittmann and McCarty (2020)[18] |
| γ-HCH | s | -86.04 | www. chemeo.com (2024)[19] |
| Chloride | aq | -131.2 | Astle et al. (1980)[20] |
| MCB | liq | 89.2 | Dolfing and Harrison (1992)[21] |
| | aq | 102.3 | |
| | g | 99.2 | |

**Table S6. Gibbs free energies of half reactions related to thermodynamic calculations in this study (pH = 7).**

| Reduced Compound | Formula | GDR* | $\Delta G^{o'}$ (kJ/e$^-$eq) | Reference |
|---|---|---|---|---|
| Ethanol | $C_2H_6O$ | 12 | 30.35 | Calculated using balanced half-reactions using standard Gibb's free energies of formation assuming bicarbonate as the oxidized form of carbon. |
| Acetate | $C_2H_3O^{2-}$ | 8 | 26.80 | |
| Hydrogen | $H_2$ | 2 | 39.87 | |
| Methane | $CH_4$ | 8 | 22.92 | |
| Lindane | $C_6H_6Cl_6$ | 24 | 57.85 | |
| MCB | $C_6H_5Cl$ | 28 | 31.46 | |
| Benzene | $C_6H_6$ | 30 | 27.37 | |
| Acetyl-CoA | $C_{23}H_{38}N_7O_{17}P_3S$ | 88 | 30.90 | McCarty (2007)[5] |

*Generalized degree of reductance (GDR) is defined as the number of electrons transferred (also referred to as the electron equivalent or e$^-$eq) during the organic half-reaction for each mole of compound. For a generic organic compound $C_nH_aO_bN_cP_dS_eCl_f$, its GDR = 4n+a-2b-3c+5d-2e-f.

**Table S7. List of half-reactions and standard Gibbs free energy used in thermodynamics calculations.**

| Reaction index | Reduced Compound | Half-reaction | $\Delta G^{0'}$ (kJ/$e^-$eq) |
|---|---|---|---|
| 1 | Lindane ($C_6H_6Cl_6$) | $\frac{1}{4}HCO_3^- + \frac{3}{2}H^+ + \frac{1}{4}Cl^- + e^- = \frac{3}{4}H_2O + \frac{1}{24}C_6H_6Cl_6$ | 57.85 |
| 2 | MCB ($C_6H_5Cl$) | $\frac{3}{14}HCO_3^- + \frac{5}{4}H^+ + \frac{1}{28}Cl^- + e^- = \frac{9}{14}H_2O + \frac{1}{28}C_6H_5Cl$ | 31.46 |
| 3 | Benzene ($C_6H_6$) | $\frac{1}{5}HCO_3^- + \frac{6}{5}H^+ + e^- = \frac{3}{5}H_2O + \frac{1}{30}C_6H_6$ | 27.37 |
| 4 | Ethanol ($C_2H_5OH$) | $\frac{1}{6}HCO_3^- + \frac{7}{6}H^+ + e^- = \frac{5}{12}H_2O + \frac{1}{12}C_2H_5OH$ | 30.35 |
| 5 | Acetate ($CH_3COO^-$) | $\frac{1}{4}HCO_3^- + \frac{9}{8}H^+ + e^- = \frac{1}{2}H_2O + \frac{1}{8}CH_3COO^-$ | 26.80 |
| 6 | Methane ($CH_4$) | $\frac{1}{8}HCO_3^- + \frac{9}{8}H^+ + e^- = \frac{3}{8}H_2O + \frac{1}{8}CH_4$ | 22.92 |
| 7 | Hydrogen ($H_2$) | $H^+ + e^- = \frac{1}{2}H_2$ | 0 |

**Table S8. Henry's law constants used in this study (measured in water at 25°C).**

| Compound | Henry's law constant, $H_{cc}$ (g/a molar concentration ratio) | Reference |
|---|---|---|
| $H_2$ | 51.1 | Morrison and Billett (1952)[22] |
| MCB | 0.127 | Shiu and MacKay (1997)[23] |
| Benzene | 0.225 | Shiu and MacKay (1997)[23] |
| Methane | 27.189 | MacKay and Shiu (1981)[24] |

**Table S9. List of notations and symbols.**

| Notation or symbol | Unit | Definition |
|---|---|---|
| AC_meth | | acetoclastic methanogenesis/methanogens |
| $b$ | day$^{-1}$ | first-order decay constant |
| $C$ | µmol/L | concentration |
| $C_s$ | mg/L | solubility limit |
| dechlr | | dechlorination |
| ferm | | fermentation |
| H2 | | hydrogen |
| H2_meth | | hydrogenotrophic methanogenesis/methanogens |
| $k$ | µmol/mg/day | maximum specific substrate utilization rate |
| $K_s$ | µmol/L | half-velocity constant |
| $K_l$ | day$^{-1}$ | first-order dissolution rate constant |
| lin | | lindane |
| meth | | methanogenesis |
| $R$ | kJ/mol/K | ideal gas constant |
| $S$ | µmol/L | substrate concentration |
| $t$ | day | time |
| $T$ | K | temperature |
| $X_a$ | mg/L | active biomass concentration |
| $Y$ | mg/µmol substrate | yield of biomass, if not specified |
| $\Psi$ | unitless | equilibrium bias as defined in Eq.2 |
| $k_{ferm}$, $k_{dechlr}$, $k_{AC\_meth}$, $k_{H2\_meth}$ | µmol/mg/day | maximum specific substrate utilization rates for ethanol fermentation, lindane dechlorination, acetoclastic methanogenesis, and hydrogenotrophic methanogenesis, respectively |
| $K_{S[ferm]}$, $K_{S\,[dechlr]}$, $K_{S\,[dechlr\_H2]}$, $K_{S\,[AC\_meth]}$, $K_{S\,[H2\_meth]}$ | µmol/L | half-velocity constants for ethanol fermentation, lindane dechlorination, hydrogen in lindane reductive dechlorination, acetoclastic methanogenesis, and hydrogenotrophic methanogenesis, respectively |
| $b_{ferm}$, $b_{dechlr}$, $b_{AC\_meth}$, $b_{H2\_meth}$ | day$^{-1}$ | decay constants for ethanol fermentation, lindane dechlorination, acetoclastic methanogenesis, and hydrogenotrophic methanogenesis, respectively |

**Table S10. Parameter estimation settings for Bayesian optimization.**

| Parameter | Unit | Range | Initial guess | Reference for initial guess |
|---|---|---|---|---|
| $k_{[EtOH,fm]}$ | μmol/(mg ·day) | 10-1000 | 720 | estimated based on Qiao et al (2019)[25] |
| $K_{S[EtOH,fm]}$ | μmol/L | 0-500 | 360 | estimated based on Qiao et al (2019)[25] |
| $k_{[HCH,dechlr]}$ | μmol/(mg ·day) | 1-200 | 20 | estimated based on Puentes Jácome (2019)[26] |
| $K_{S[HCH,dechlr]}$ | μmol/L | 0-500 | 210 | estimated based on Puentes Jácome (2019)[26] |
| $K_{S[H_2,dechlr]}$ | μmol/L | 0-1 | 0.1 | Fennell (1998)[27] |
| $k_{[H_2,meth]}$ | μmol/(mg ·day) | 10-1000 | 163 | Bagley (1998)[28] |
| $K_{S[H_2,meth]}$ | μmol/L | 0-1 | 0.96 | Bagley (1998)[28] |
| $k_{[AC,meth]}$ | μmol/(mg ·day) | 1-200 | 56 | Bagley (1998)[28] |
| $K_{S[AC,meth]}$ | μmol/L | 0-1000 | 450 | Bagley (1998)[28] |
| $b_{[EtOH,fm]}$ | $day^{-1}$ | 0-0.05 | 0.05 | Rittman and McCarty (2012)[29] |
| $b_{[HCH,dechlr]}$ | $day^{-1}$ | 0-0.05 | 0.05 | Rittman and McCarty (2012)[29] |
| $b_{[H_2,meth]}$ | $day^{-1}$ | 0-0.05 | 0.01 | Rittman and McCarty (2012)[29] |
| $b_{[AC,meth]}$ | $day^{-1}$ | 0-0.05 | 0.01 | Rittman and McCarty (2012)[29] |

**Tables S11-S16 are large data tables in a separate Excel file.**

**Table S11. qPCR and related data for decay constant estimation.**

**Table S12. Raw data of chemical measurements in the biokinetics experiment.**

**Table S13. Calibration dataset from bottle EXP2 as well as model initialization data for parameter estimation.**

**Table S14. Metadata of next generation sequencing in the biokinetics experiment.**

**Table S15. Raw reads of bacterial amplicon sequencing variants from next generation sequencing.**

**Table S16. Raw reads of archaeal amplicon sequencing variants from next generation sequencing.**

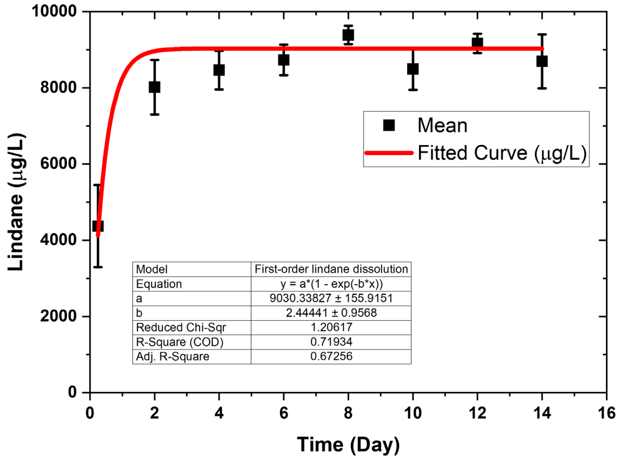

**Figure S1. First-order lindane dissolution kinetics modelled using lindane concentration data from the analytical control group bottles (AC1, AC2, and AC3) during the first 14 days. Error bars denote group standard deviations.**

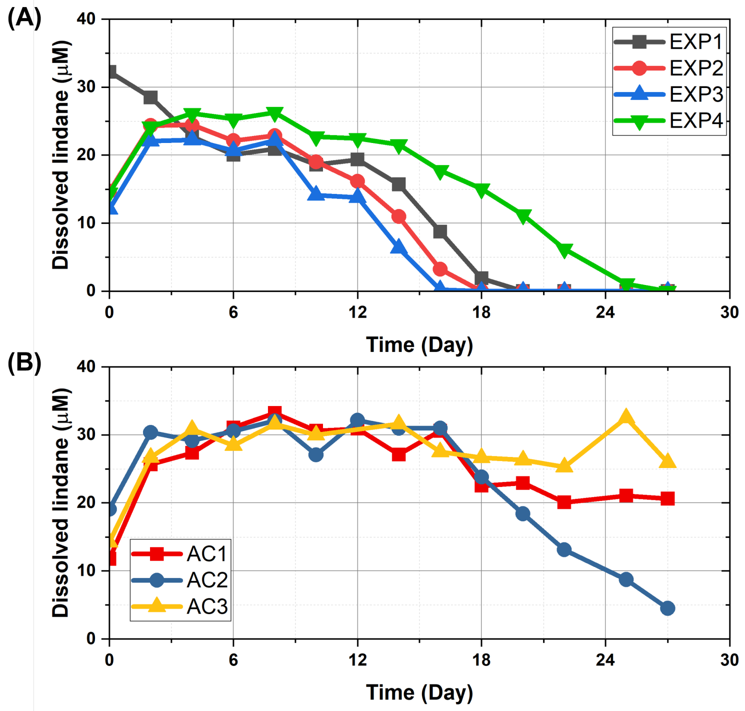

**Figure S2. Dissolved lindane concentrations measured in individual bottles from (A) the experimental group (EXP1-EXP4) and (B) the analytical control group (AC1-AC3).**

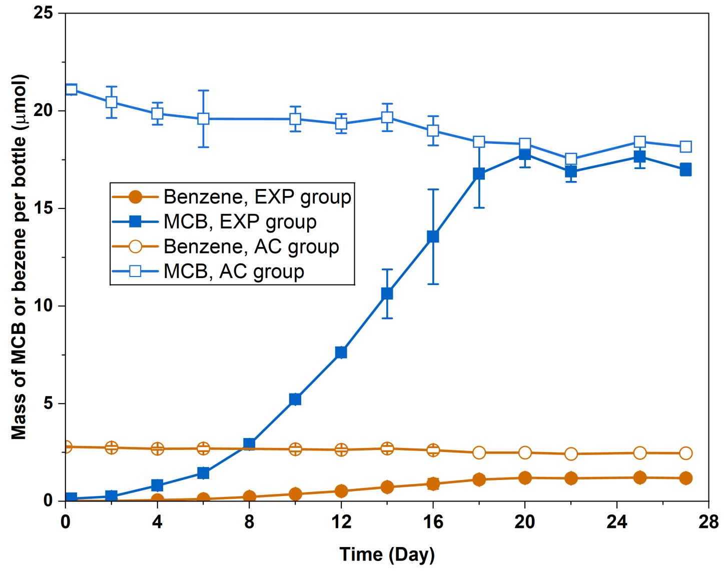

**Figure S3. Masses of monochlorobenzene (MCB; squares) and benzene (circles) per bottle in the experimental group (solid symbols) and the analytical control group (open symbols), error bars denote within-group standard deviations.**

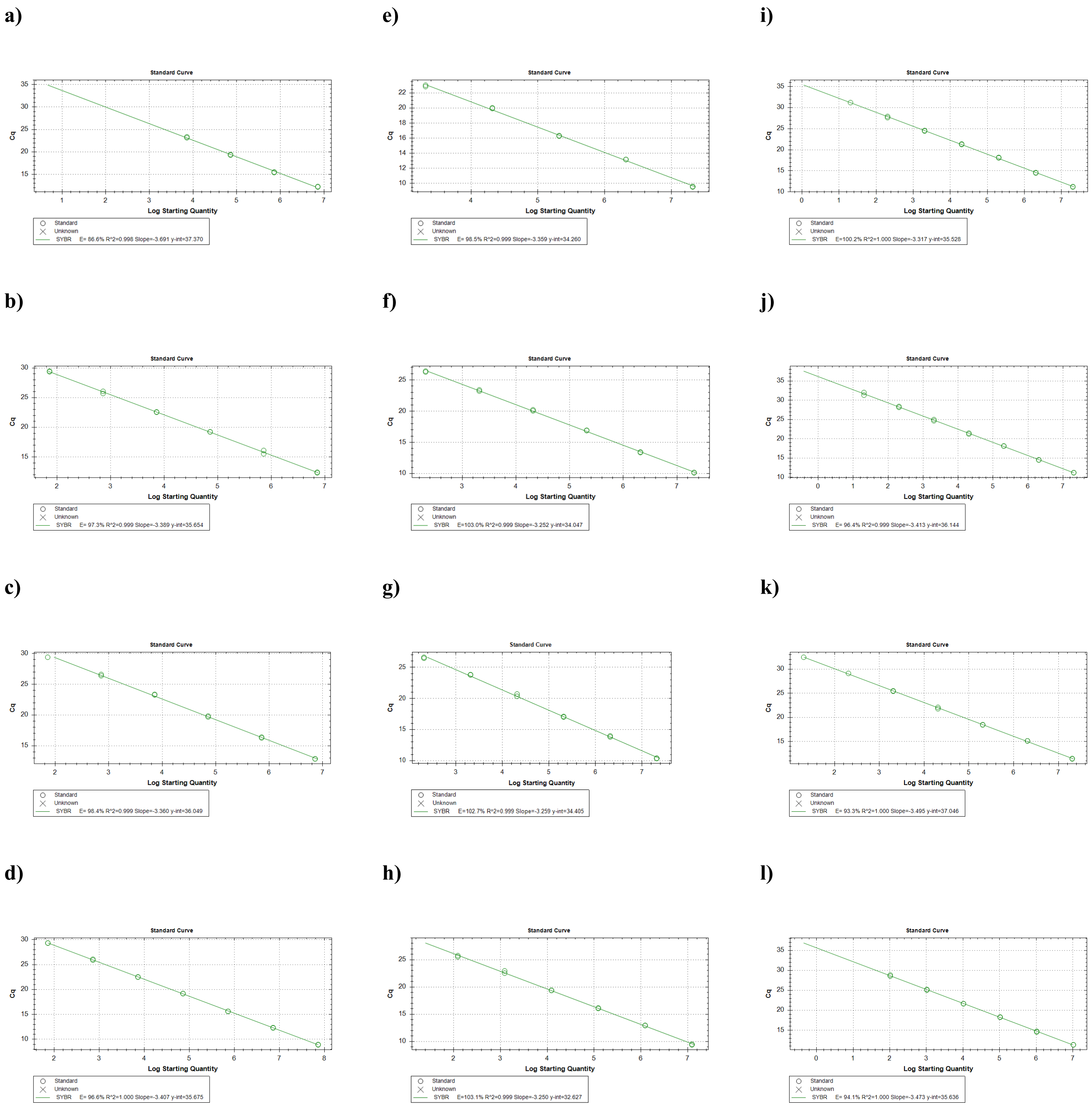

**Figure S4. Standard calibration curves and amplification efficiencies of all qPCR assays carried out in this study for quantifying a)-d) *Dehalobacter*, e)-h) total bacteria (GenBac), and i)-l), total archaea (GenArch).**

(A)

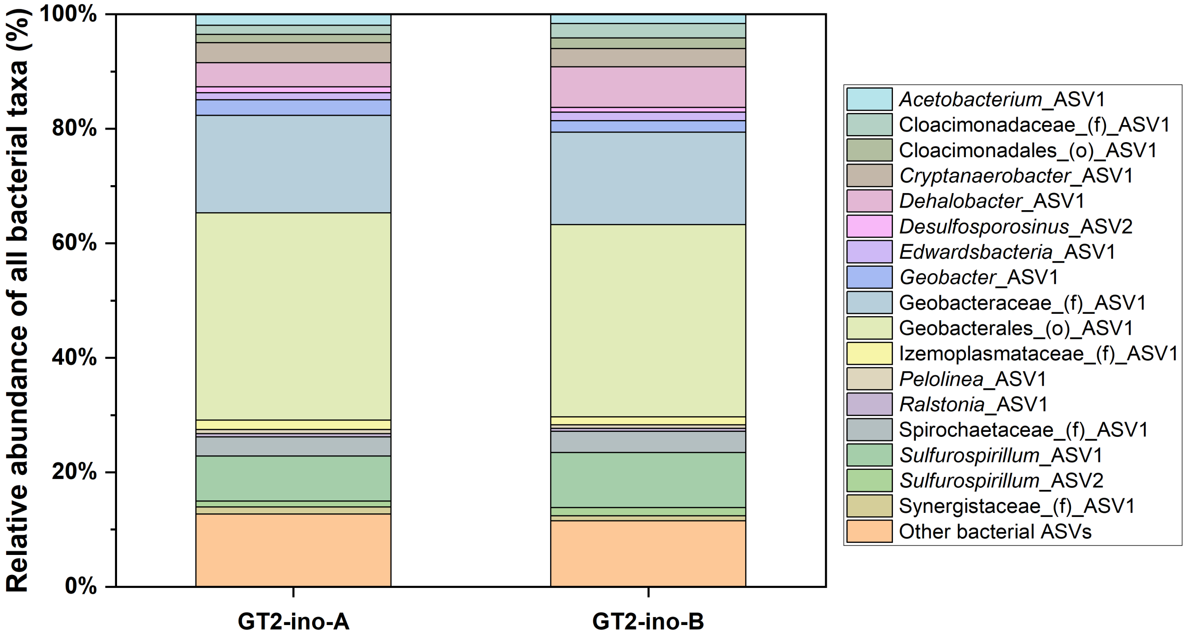

(B)

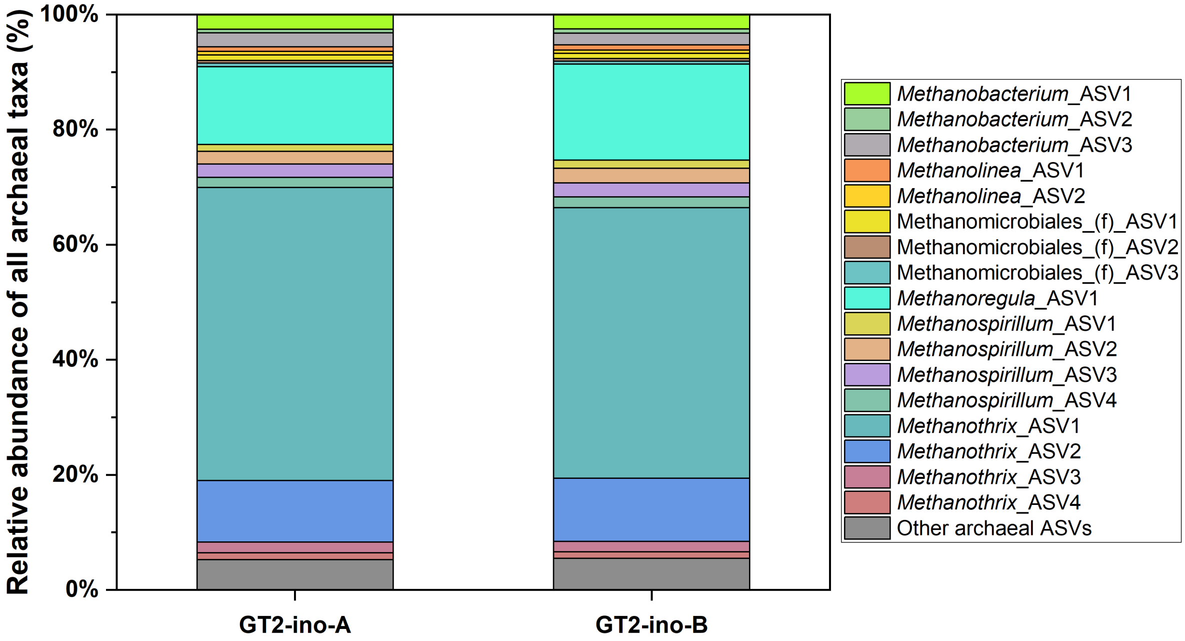

**Figure S5. Microbial community structure at the ASV level of replicate DNA samples of the GT2 inoculum culture used for the batch experiment: (A) Bacteria, (B) Archaea.**

(A)

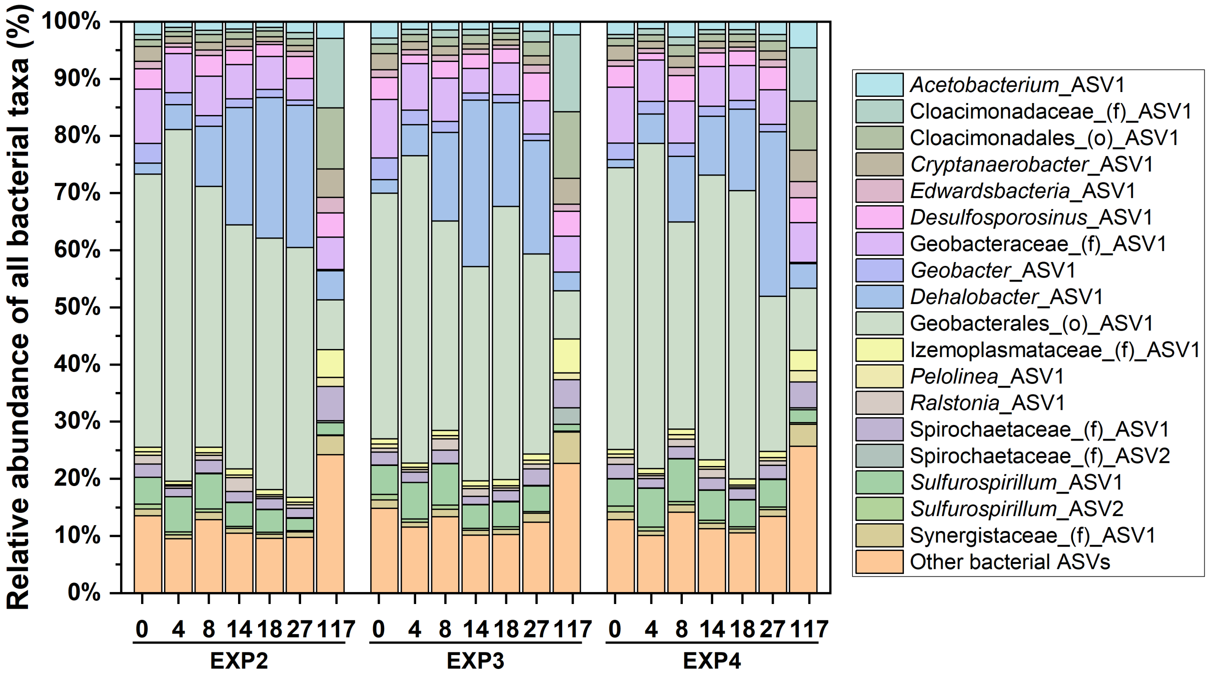

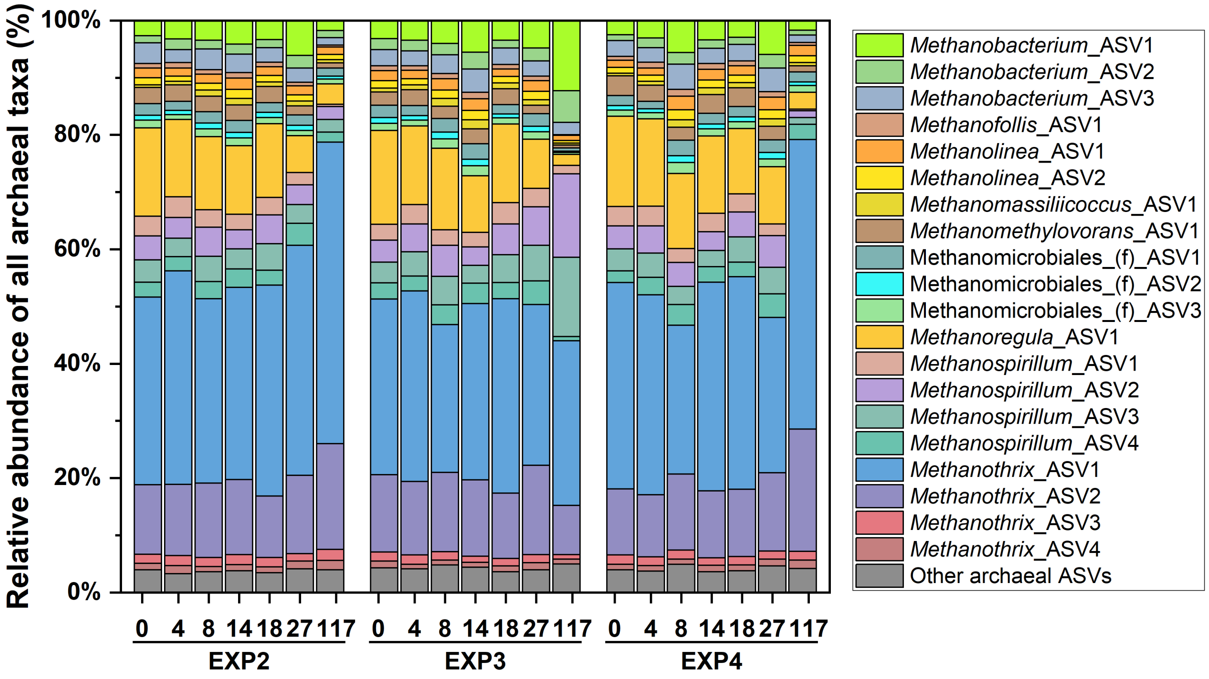

(B)

**Figure S6. Microbial community structure at the amplicon sequence variants (ASV) level for bottles EXP2, EXP3 and EXP4 of the experimental (EXP) group: (A) Bacteria, (B) Archaea.**

(A)

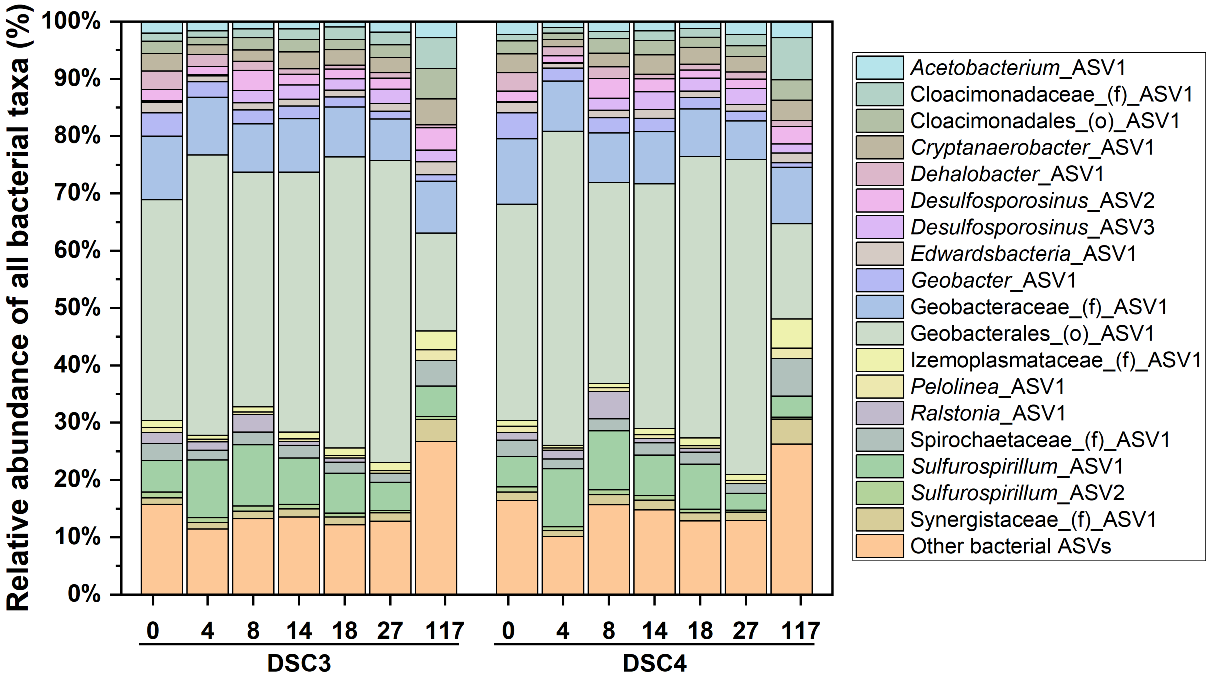

(B)

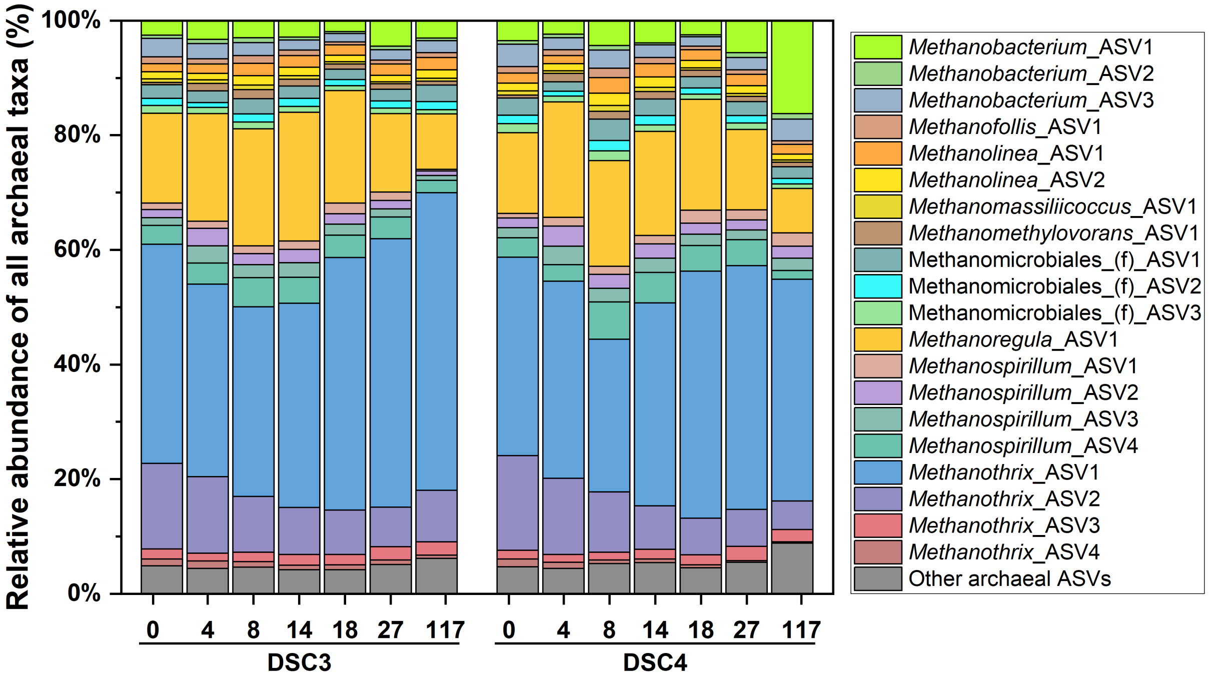

**Figure S7. Microbial community structure at the amplicon sequence variants (ASV) level for bottles DSC3 and DSC4 of the *Dehalobacter* starved control (DSC) group: (A) Bacteria, (B) Archaea.**

(A)

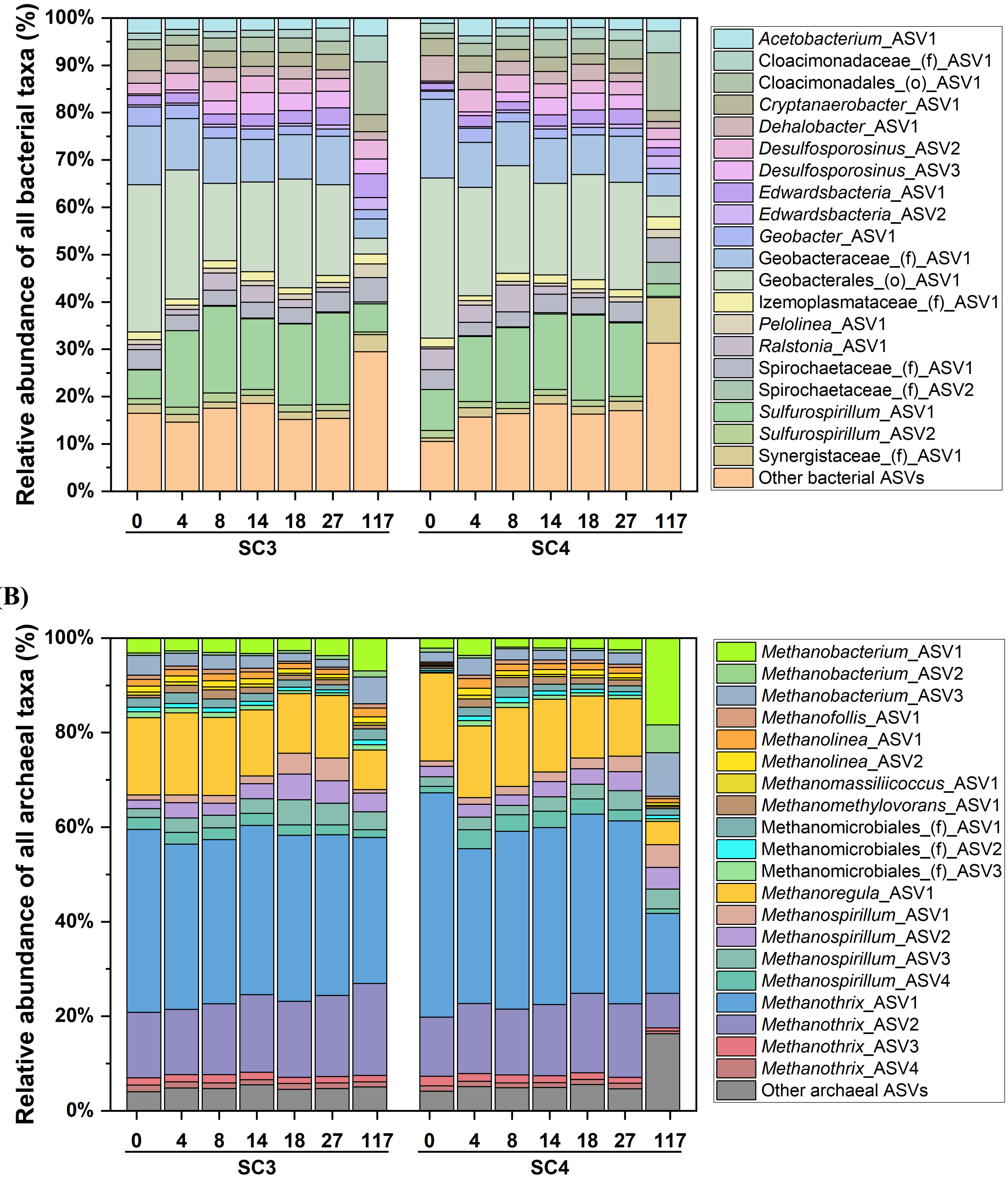

**Figure S8. Microbial community structure at the amplicon sequence variants (ASV)level for bottles SC3 and SC4 of the starved control (SC) group: (A) Bacteria, (B) Archaea.**

(A)

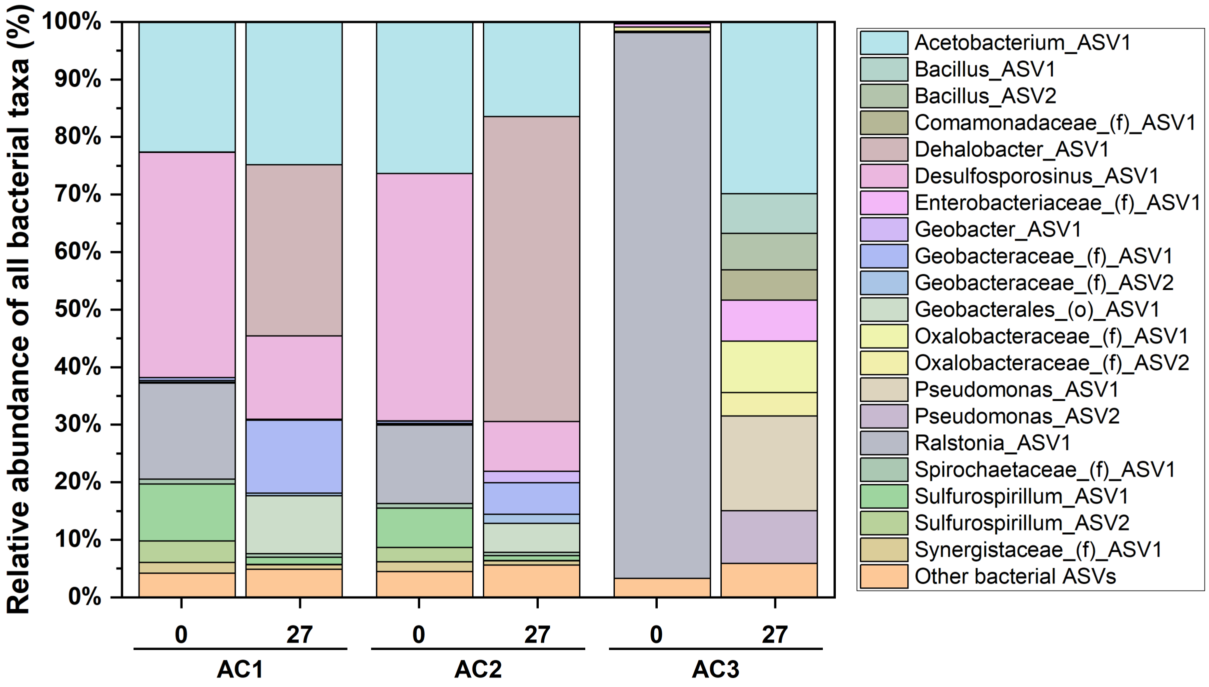

(B)

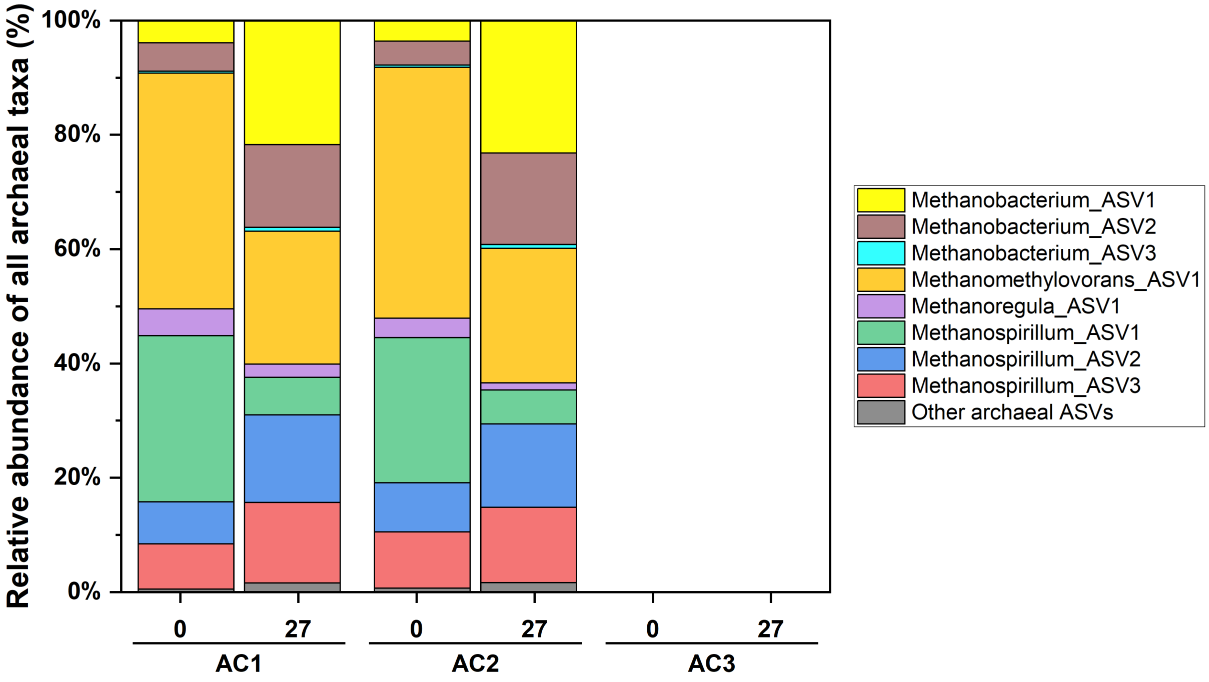

**Figure S9. Microbial community structure at the amplicon sequence variants (ASV) level of analytical control (AC) group bottles: (A) Bacteria, (B) Archaea. Note: archaeal DNA was not detected in bottle AC3.**

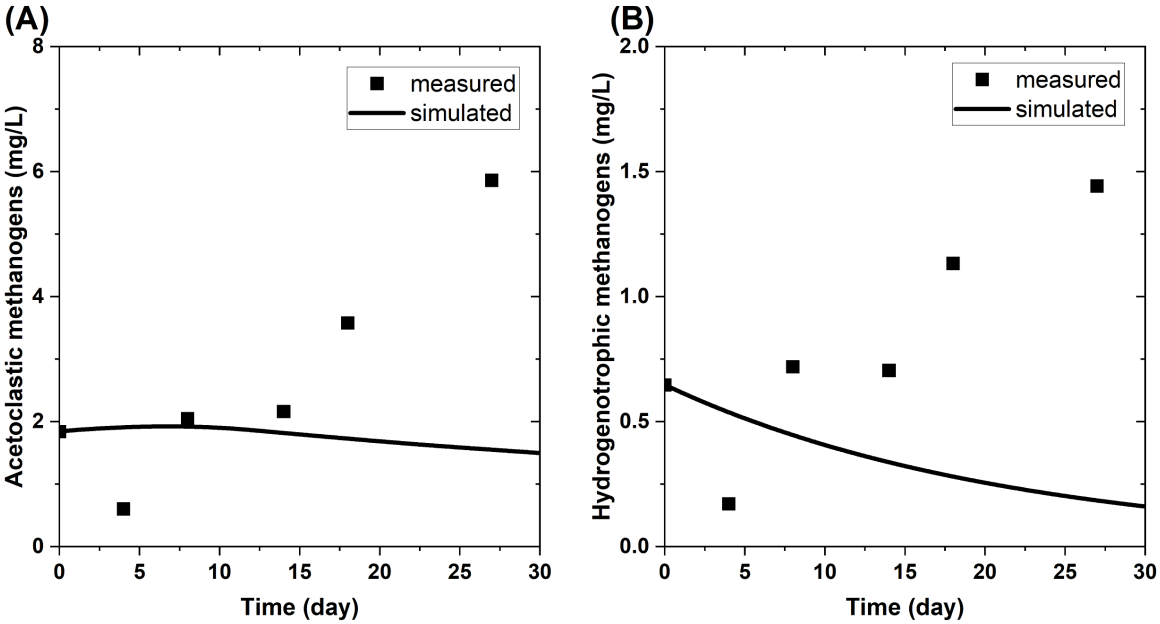

**Figure S10. Measured and simulated methanogens biomass data. A: acetoclastic methanogens, B: hydrogenotrophic methanogens. Time range used for fitting for each data series correspond to those used in parameter estimation.**

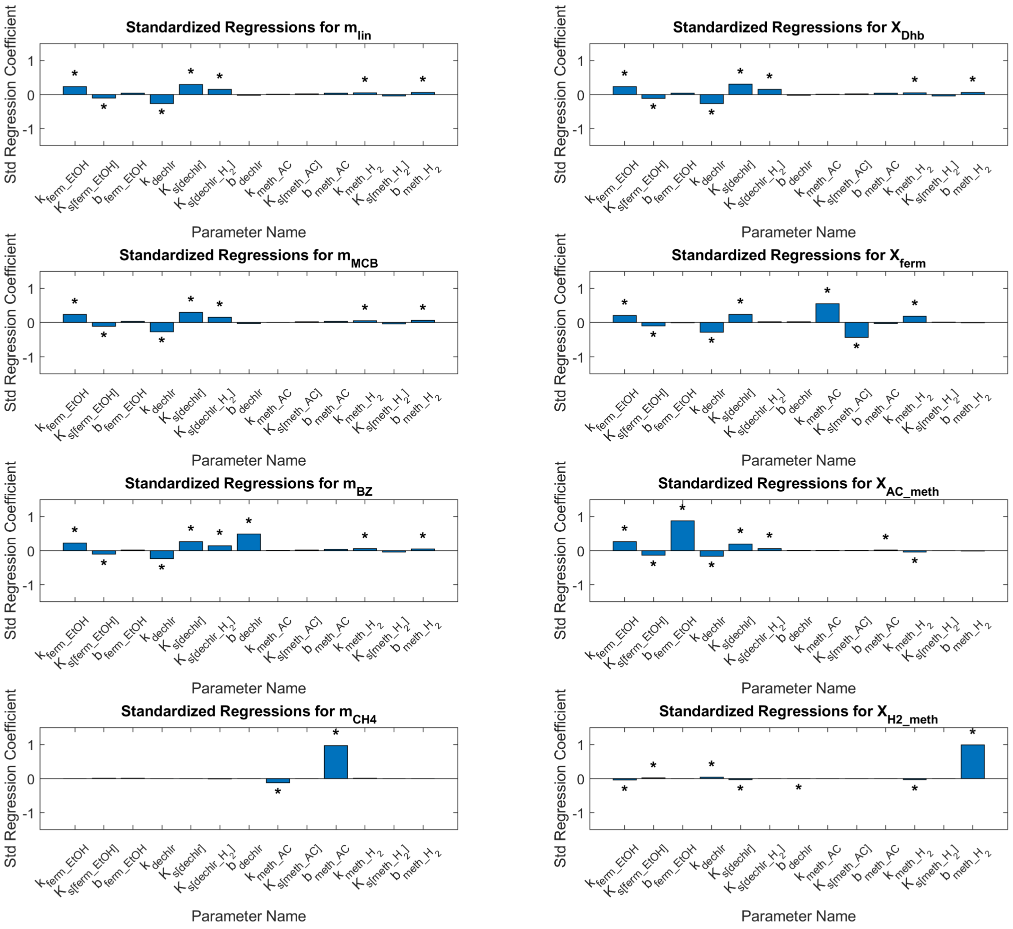

**Figure S11. Standardized regression analysis of individual model output, including mass of lindane ($m_{lin}$), MCB ($m_{MCB}$), benzene ($m_{BZ}$), methane ($m_{CH4}$), and biomass concentrations of *Dehalobacter* ($X_{Dhb}$), ethanol fermenter ($X_{ferm}$), acetoclastic methanogens ($X_{AC\_meth}$), and hydrogenotrophic methanogens ($X_{H2\_meth}$). Asterisks mark parameters that are statistically significant at the 0.05 level.**

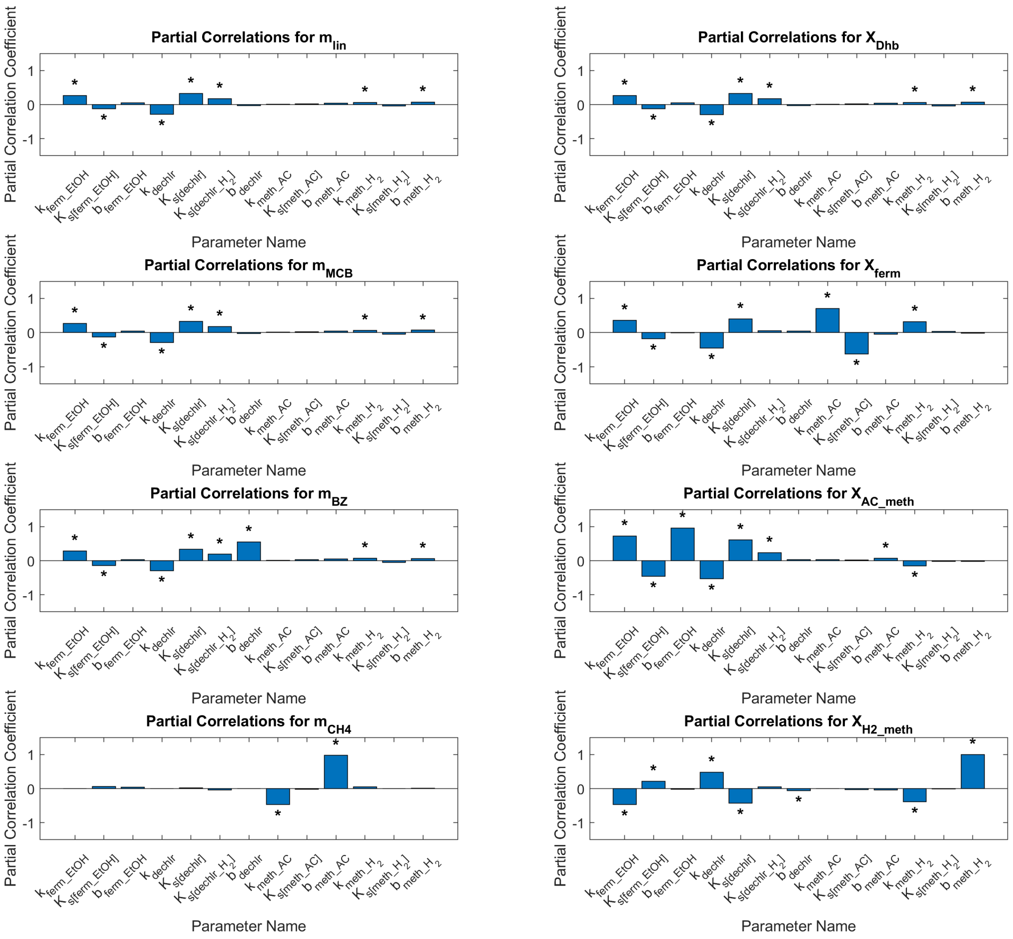

**Figure S12. Partial correlation analysis of individual model output, including mass of lindane ($m_{lin}$), MCB ($m_{MCB}$), benzene ($m_{BZ}$), methane ($m_{CH4}$), and biomass concentrations of *Dehalobacter* ($X_{Dhb}$), ethanol fermenter ($X_{ferm}$), acetoclastic methanogens ($X_{AC\_meth}$), and hydrogenotrophic methanogens ($X_{H2\_meth}$). Asterisks mark parameters that are statistically significant at the 0.05 level.**

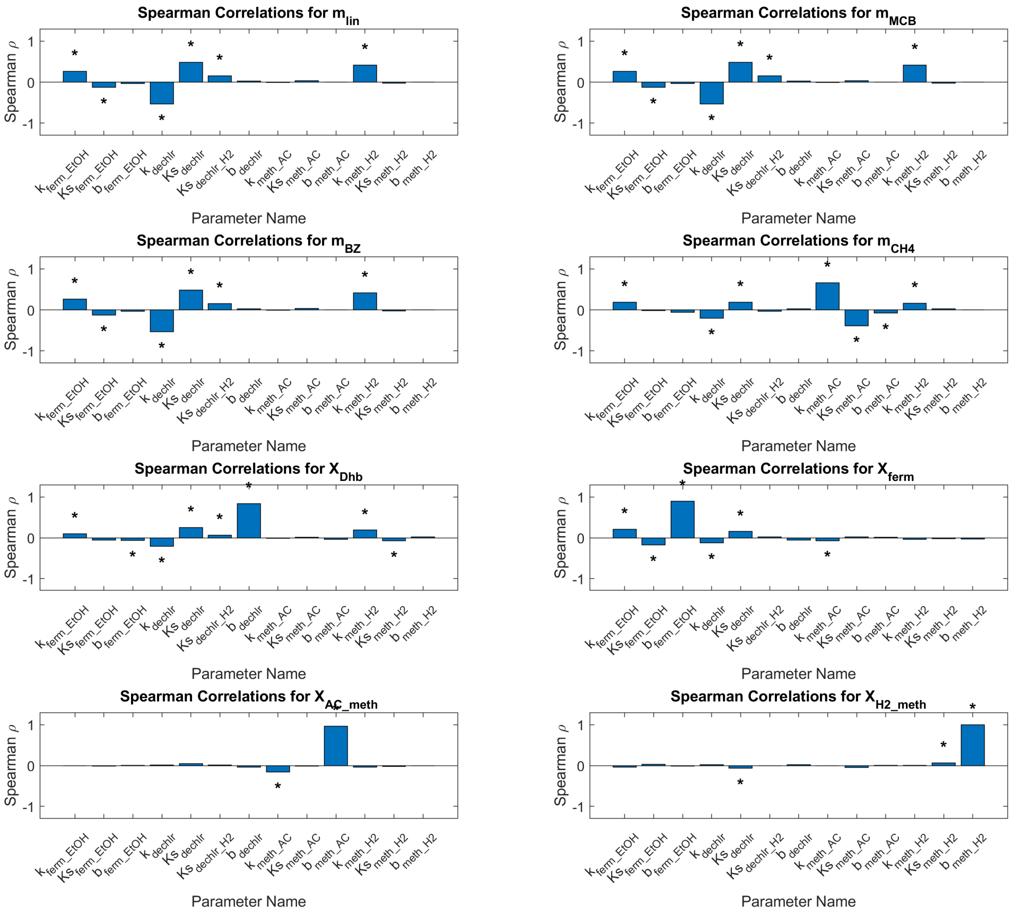

**Figure S13. Spearman correlation analysis of individual model output, including mass of lindane ($m_{lin}$), MCB ($m_{MCB}$), benzene ($m_{BZ}$), methane ($m_{CH4}$), and biomass concentrations of *Dehalobacter* ($X_{Dhb}$), ethanol fermenter ($X_{ferm}$), acetoclastic methanogens ($X_{AC\_meth}$), and hydrogenotrophic methanogens ($X_{H2\_meth}$). Asterisks mark parameters that are statistically significant at the 0.05 level.**

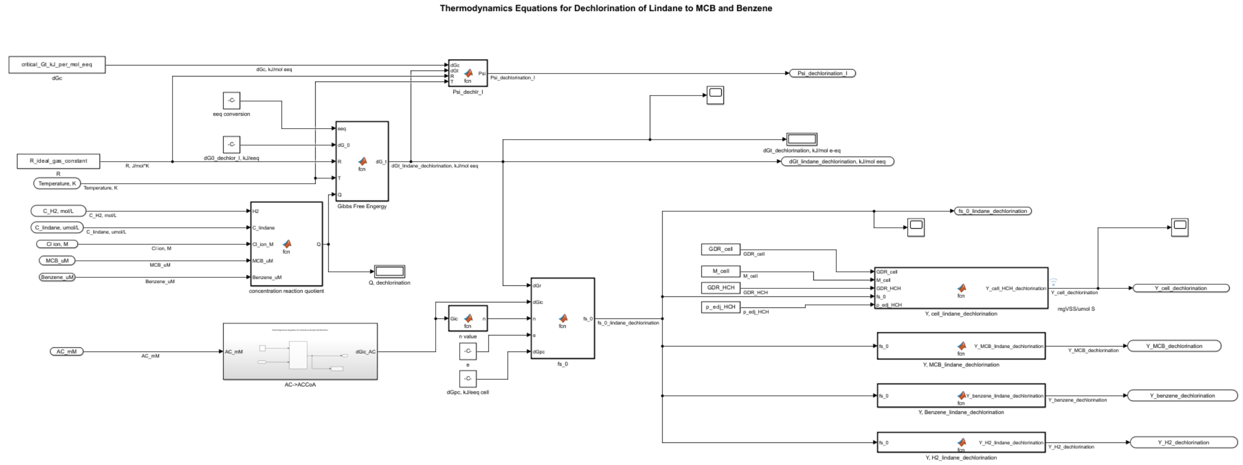

**Figure S14. Example Simulink model screenshot: thermodynamics block for lindane dechlorination. Signal flow follows the direction of the connected arrows. Regular and rounded rectangular symbols denote input or output scalar signals, symbols with the MATLAB icon denote coding blocks, and scope symbols denote signal-monitoring blocks for tracking output values.**

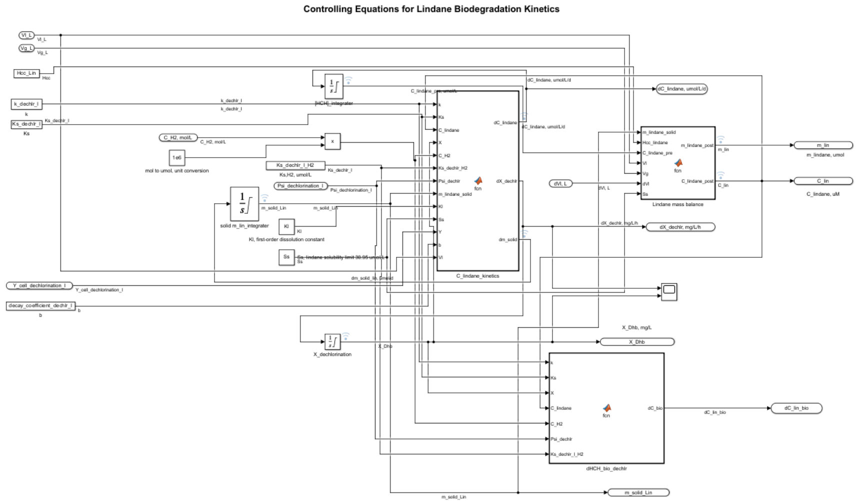

**Figure S15. Example Simulink model screenshot: biokinetics block for lindane dechlorination. Signal flow follows the direction of the connected arrows. Regular and rounded rectangular symbols denote input or output scalar signals, symbols with the**

**MATLAB icon denote coding blocks, scope symbols denote signal-monitoring blocks for tracking output values, and symbols with integrator signs denote integration of input signals.**

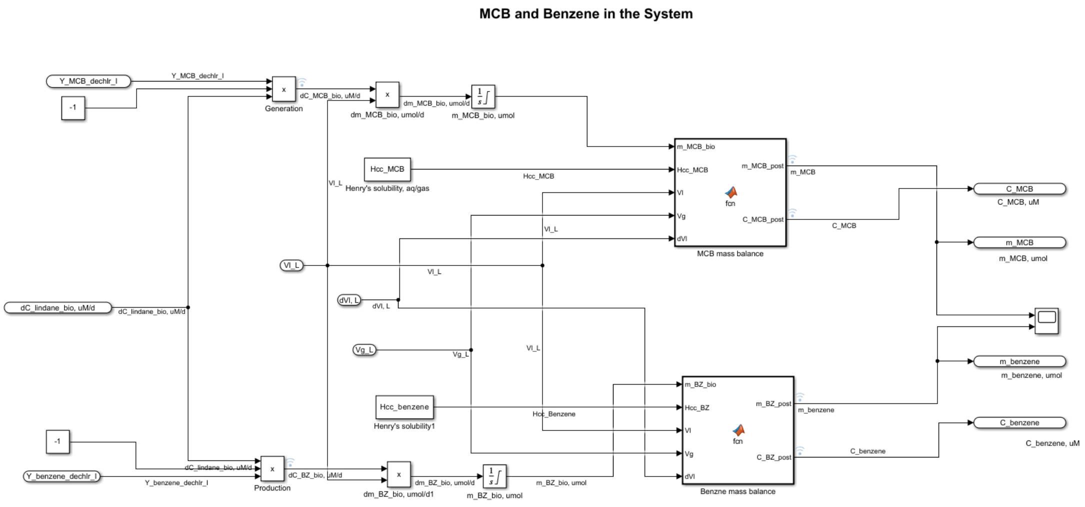

**Figure S16. Example Simulink model screenshot: mass balance block for MCB and benzene. Signal flow follows the direction of the connected arrows. Regular and rounded rectangular symbols denote input or output scalar signals, symbols with the MATLAB icon denote coding blocks, scope symbols denote signal-monitoring blocks for tracking output values, symbols with integrator signs denote integration of input signals, and symbols with multiplication signs denote multiplication of input signals.**
